# Temporal patterns in *Ixodes ricinus* microbial communities: an insight into tick-borne microbe interactions

**DOI:** 10.1101/2020.09.26.314179

**Authors:** E Lejal, J Chiquet, J Aubert, S Robin, A Estrada-Peña, O Rue, C Midoux, M Mariadassou, X Bailly, A Cougoul, P Gasqui, JF Cosson, K Chalvet-Monfray, M Vayssier-Taussat, T Pollet

**Author notes:** **Corresponding author:** Dr Thomas Pollet.

## Abstract

**Background:** Ticks transmit pathogens of medical and veterinary importance, and represent an increasing threat for human and animal health. Important steps in assessing disease risk and developing possible new future control strategies involve identifying tick-borne microbes, their temporal dynamics and interactions.

**Methods:** Using high throughput sequencing, we studied the microbiota dynamics of *Ixodes ricinus* from 371 nymphs collected monthly over three consecutive years in a peri-urban forest. After adjusting a Poisson Log Normal model to our data set, the implementation of a principal component analysis as well as sparse network reconstruction and differential analysis allowed us to assess inter-annual, seasonal and monthly variability of *I. ricinus* microbial communities as well as their interactions.

**Results:** Around 75% of the detected sequences belonged to five genera known to be maternally inherited bacteria in arthropods and potentially circulating in ticks: Candidatus *Midichloria, Rickettsia, Spiroplasma, Arsenophonus* and *Wolbachia*. The structure of the *I. ricinus* microbiota was temporally variable with interannual recurrence and seemed to be mainly driven by OTUs belonging to environmental genera. The total network analysis revealed a majority of positive (partial) correlations. We identified strong relationships between OTUs belonging to *Wolbachia* and *Arsenophonus*, betraying the presence of the parasitoid wasp *Ixodiphagus hookeri* in ticks, and the well known arthropod symbiont *Spiroplasma*, previously documented to be involved in the defense against parasitoid wasp in *Drosophila melanogaster*. Other associations were observed between the tick symbiont Candidatus *Midichloria* and pathogens belonging to *Rickettsia*, probably *Rickettsia helvetica*. More specific network analysis finally suggested that the presence of pathogens belonging to genera *Borrelia, Anaplasma* and *Rickettsia* might disrupt microbial interactions in *I. ricinus*.

**Conclusions:** Here, we identified the *I. ricinus* microbiota and documented for the first time the existence and recurrence of marked temporal shifts in the tick microbial community dynamics. We statistically showed strong relationships between the presence of some pathogens and the structure of the *I. ricinus* non-pathogenic microbes. We interestingly detected close links between some tick symbionts and the potential presence of either pathogenic *Rickettsia* or a parasitoid in ticks. All these new findings might be very promising for the future development of new control strategies of ticks and tick-borne diseases.

## Introduction

Ticks, vectors of several zoonotic pathogens, represent an important and increasing threat for human and veterinary health. While these arthropods are among the most important vectors of pathogens affecting humans and animals worldwide, it is now well established that tick-borne pathogens (TBPs) coexist with many other microorganisms in ticks. These other tick-borne microbes, commensal and/or symbionts, are likely to confer multiple detrimental, neutral, or beneficial effects to their tick hosts, and can play various roles in nutritional adaptation, development, reproduction, defense against environmental stress, and immunity [1,2]. Otherwise, they may also interact with tick-borne pathogens and thus influence the tick vector competence [3,4]. In this context, the identification and characterization of tick microbiota has become crucial to better understand the tick-microbe interactions. With the development of high throughput sequencing technologies, the number of studies dealing with tick microbiota considerably increased in the past ten years and revealed an unexpected microbial diversity in ticks [5–14]. The microbiota of several tick species of the genera *Ixodes, Dermacentor, Haemaphysalis, Rhipicephalus* and *Amblyomma* has been studied (see in [15]) and all these works have given crucial details on microbial communities in ticks and improved our knowledge on tick microbial community ecology. However, few information are currently available on the microbiota composition of the tick *Ixodes ricinus* [6,12,16–22], the predominant tick species in western Europe, able to transmit the largest variety of pathogens, including the Lyme disease agent *Borrelia burgdorferi* s.l.. Moreover, several of these previous *I. ricinus* microbiota analyses may have overestimated the microbiota diversity due to extraction step contamination and the lack of controls in their analysis [22]. Studies dealing with *I. ricinus* microbial communities pointed out complex microbial assemblages inhabiting ticks as pathogens, specific endosymbionts, commensal and environmental microbes. As already observed for pathogens [23–26], the other members of these biological assemblages are probably dynamic and likely to vary with time. While the scale affects our view on the dynamics of tick microbial communities [27], few information are currently available on the temporal patterns of the *I. ricinus* microbiota. Does the *I. ricinus* microbiota vary with season? Are these potential temporal patterns annually redundant? Answering these questions is a crucial first step to keep better understanding both the tick and tick-microbe ecology and identify. Otherwise, while it is now admitted that tick microbiota may also play a role in driving transmission or multiplication of tick-borne pathogens [28], no information is currently available on the interactions between *I. ricinus-borne* microbes and on potential co-occurrences between the presence of pathogens and the *I. ricinus* microbiota. These information are essential to identify potential new control strategies of ticks and tick-borne diseases in the future. After monthly collecting *I. ricinus* nymphs over three consecutive years in a peri-urban forest, we used a high throughput sequencing approach to identify the *I. ricinus* microbiota and assess its temporal dynamics. Thanks to multivariate and partial correlation network analyses, we identified direct statistical associations between members of the microbiota, including pathogenic genera, and assessed the influence of TBPs presence on tick microbiota structure and interactions.

## Material and Methods

### Tick collection

Questing *Ixodes ricinus* nymphs were collected for three years by dragging (from April 2014 to May 2017) in the Sénart forest in the south of Paris. More details on the sampling location and design, and tick collection, are available in Lejal *et al*. [26].

### Tick homogenisation and DNA extraction

998 nymphs have been collected over the three years [26]. As detailed in our previous studies [22,26,29], ticks were first washed once in ethanol 70% for 5 minutes and rinsed twice in sterile MilliQ water. They were then individually homogenised in 375 μL of Dulbecco’s Modified Eagle Medium with decomplemented foetal calf serum (10%) and six steel beads using the homogenizer Precellys^®^24 Dual (Bertin, France) at 5,500 rpm for 20 seconds. DNA extraction was performed on 100 μL of tick homogenate, using the NucleoSpin^®^ Tissue DNA extraction kit (Macherey-Nagel, Germany).

### DNA amplification and multiplexing

Among the 998 nymphs, 557 have been chosen to characterize the temporal dynamics. Thanks to our previous study [26], we knew the infection rate of all collected ticks. For each collecting month, while all infected nymphs have been compulsory integrated in the analysis, non-infected nymphs (at least 15 per month) were added until reaching a minimum of 30 ticks (infected and non-infected) per month, when it was possible. When less than 30 ticks were collected in a month, all ticks were added in the analysis. In addition to tick samples, 45 negative controls have been performed to distinguish tick microbial OTUs from contaminants [22]. As detailed in Lejal *et al*. [22], DNA amplifications were performed on the V4 region of the 16s rRNA gene using the primer pair used by Galan *et al*. [30] (16S-V4F: 5’-GTGCCAGCMGCCGCGGTAA-3’ and 16S-V4R: 5’-GGACTACHVGGGTWTCTAATCC-3’), producing a 251 bp amplicon. Different 8bp indexes were added to primers allowing *in fine* the amplification and multiplexing of all samples. All the PCR amplifications were carried out using the Phusion^®^ High-Fidelity DNA Polymerase amplification kit (Thermo Scientific, Lithuania). For each sample, 5 μL of DNA extract were amplified in a 50 μL final reaction volume, containing 1X Phusion HF buffer, 0.2 μM of dNTPs, 0.2 U/mL of Phusion DNA polymerase, and 0.35 μM of forward and reverse primer. The following thermal cycling procedure was used: initial denaturation at 98°C for 30 seconds, 35 cycles of denaturation at 98°C for 10 seconds, annealing at 55°C for 30 seconds, followed by extension at 72°C for 30 seconds. The final extension was carried out at 72°C for 10 minutes. PCR products were checked on 1.5% agarose gels, cleaned and quantified and finally pooled at equimolar concentrations and sent to the sequencing platform (GenoScreen, France) [22].

### Sequencing and data processing

The equimolar mix was concentrated and sequenced by GenoScreen (Lille, France) using MiSeq Illumina 2 x 250 bp chemistry. All the quality controls and different steps of sequence analyses have been performed (more details are available in [22]). Based on results obtained in the 45 negative controls, sequences considered as contaminants have been removed from the dataset [22]. Due to this large OTUs filtration, we identified several samples with less than 500 sequences. We considered this number of sequences too low to be analysed and we thus removed the concerned samples. Finally, the microbiota of 371 *I. ricinus* nymphs was analysed from a final dataset composed of 907,941 sequences.

### Data preprocessing

Three distinct datasets have been analysed during this study. First, the entire one, including all the nymphs analysed in 16s rRNA gene sequencing, except the only two nymphs collected in November (369 analysed samples). Second, the No *Wolbachia/Arsenophonus* dataset, corresponding to a reduced dataset, whose samples harboring OTUs belonging to *Arsenophonus* and/or *Wolbachia* genera (presenting a number of sequences significantly higher than the highest number of sequences detected for the same OTUs in negative controls, in a 95% confidence interval) have been removed (303 samples analysed in this data set). Third, the TBPs dataset, composed of samples undoubtedly identified as TBPs positive for only one genus of TBPs (co-infected samples were not included due to the too low number of samples and the difficulty to interpret the results) according to 16s rRNA gene sequencing results as well as microfluidic PCR detection [26]. We considered a sample as positive if a TBP species was previously detected in microfluidic PCR and if at least one OTUs identified in 16s rRNA gene sequencing, corresponding to the same pathogenic genera, presented a number of sequences significantly higher than the highest number of sequences detected for this OTUs in negative controls (in a 95% confidence interval). Before analyzing the data, we applied standard filtering to all the three data sets to remove OTUs associated with weak counts that could hamper the statistical analysis. First, OTUs whose total number of sequences was lower than the total number of samples have been removed. Second, a filter related to the yearly prevalence of each OTU: those consistently detected in less than 10% of the samples of each year have been removed. The application of these filters on the three datasets, the entire, the No *Wolbachia/Arsenophonus* dataset and the TBPs dataset, lead to the selection of 89 and 82 and 74 OTUs to be included in the subsequent statistical analyses, respectively.

### Statistical analysis

The statistical framework used to describe our data sets is the multivariate Poisson lognormal (PLN) distribution [31]. This statistical distribution is adapted to multivariate count data and shows an expected over-dispersion property compared to the standard Poisson distribution. The main idea behind the PLN model is to represent all the dependency structure between the OTUs in a latent (hidden) multivariate Gaussian layer, while a Poisson distribution in the observation space of the data is used to model counts and noise. Moreover, the PLN framework can be easily interpreted as a Generalized Multivariate Linear Model (GLM); thus it naturally allows one to include effects of covariates or offsets like in a standard GLM. All the statistical analyses performed in our paper rely on the PLN model and the variants implemented in the R package PLNmodels (version 0.11.0-9005) [32]. In particular, PLNmodels includes variants to perform standard multivariate analyses for count tables, such as PCA (Principal Component Analysis), LDA (Linear Discriminant Analysis), or sparse network reconstruction (aka sparse inference of (inverse) covariance matrix). Additional methodological details can be found in Chiquet *et al*. [33,34].

In all the models fitted in this paper, we accounted for the sampling effort (that is, the sequencing depth of each sample) by adding an offset term corresponding to the (log) total sum of count per sample obtained prior to any filtering. Moreover, in order to correct for any spurious effect induced by a given year of sampling that may hide others effects such as seasonality, we included a covariate to account for the year of sampling in the two first models implying the use of both the entire and the No *Wolbachia/Arsenophonus* dataset. While investigating the effect of TBPs on tick microbiota, we also systematically included covariates to account for the year and the season of sampling while dealing with the TBPs dataset. A covariate accounting for the presence of TBPs was also considered in the establishment of TBPs network in order to evaluate the importance of this variable on tick microbiota network establishment.

Principal component analysis was performed with the PCA variant of PLN and the PLNPCA R function [34], which performs probabilistic Poisson PCA. Network analysis was performed with the PLNnetwork function, which adds a sparsity constraint on the inverse covariance matrix in the latent Gaussian layer [33].

A differential analysis was also performed using the edgeR package (version 3.30.0) [35] on the TBPs dataset to compare OTUs abundances between positive and negative samples. *edgeR* uses the negative binomial (NB) distribution to model the read counts for each OTU in each sample and compute an empirical Bayes estimate of the NB dispersion parameter for each OTU, with abundance levels are specified by a log-linear model. For the same reasons as previously described, we introduced two covariates for the year and the season in addition to the group main effect (TBP positive or negative). Data were normalized for differences between library sizes using-the Trimmed Mean of M value (TMM) method [36]. Models are fitted with the edgeR::glmFit function, which implements generalized linear methods developed by McCarthy *et al*. [37]. For each OTU, we tested the group effect using likelihood-ratio statistics (edgeR::glmLRT function). Differentially abundant OTUs were defined as those with a p-values <0.05 after adjustment for multiple testing using the Bonferroni procedure.

Networks generated from the TBPs dataset were also compared to each other. For this purpose, a weighted version of Kendall’s τ, which integrates the edge appearance rank within families of networks (negative, *Rickettsia, Borrelia, Anaplasma* and total corrected for TBPs effects), was calculated and used to compare them with each other.

All the statistical analyses were performed on R 4.0.2. [38].

## Results

### • *Ixodes ricinus* microbiota diversity and composition

Considering the microbiota of all the 371 nymphs, we detected 353 OTUs. 307 belonged to 109 identified genera, and 46 OTUs belonged to multi-affiliated or unknown genera spread in 15 families. The mean Shannon diversity index was previously estimated to 2.1 (SD= +/- 0.8) and varied between 0.3 (nymph collected in April 2016) and 3.8 (nymph collected in October 2014) [22]. Bacterial genera with proportions higher than 0.5% of all sequences in the dataset belonged to *Arsenophonus*, Candidatus *Midichloria, Rickettsia, Wolbachia, Spiroplasma, Methylobacterium, Mycobacterium, Pseudomonas, Stenotrophomonas, Williamsa, Rickettsiella, Chryseobacterium, Borrelia, Bacillus, Anaplasma, Allorhizobium-Neorhizobium-Pararhizobium-Rhizobium* and two multi-affiliated OTUs belonging to Rhizobiaceae and Microbacteriaceae families. In total, these sequences represented 93% of all sequences in the dataset (**Figure 1**).

**Figure 1:**
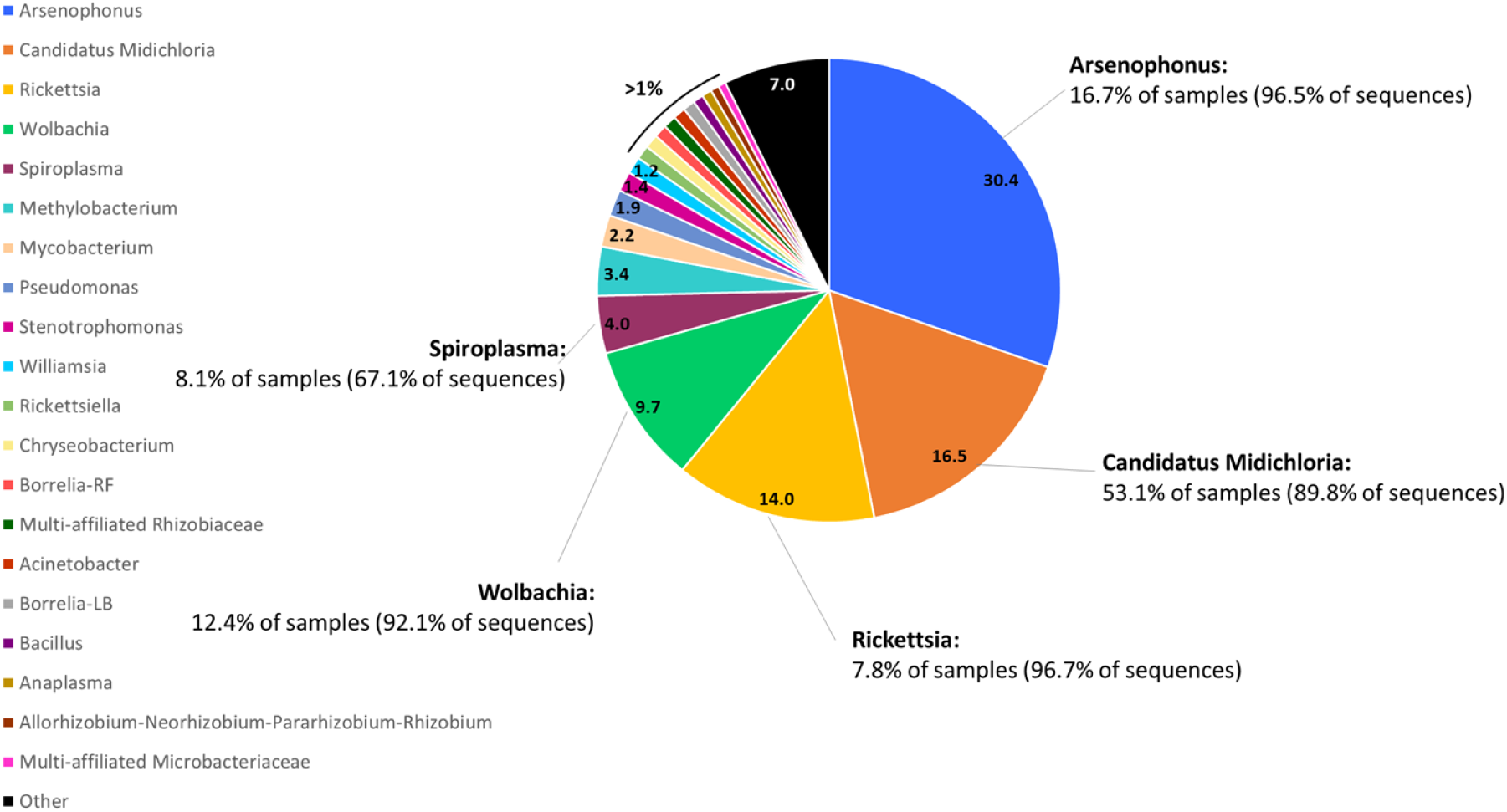
Most dominant genera in the *Ixodes ricinus* microbiota. Selected genera and multi-affiliated OTUs are those representing more than 0.5% of the total number of sequences detected in the entire dataset. Numbers given in the pie chart correspond to this percentage.

Although these genera represent a large number of sequences identified in the dataset, it is important to note that their presence is not equally distributed in all the samples. To investigate this point, we determined, for the five first represented genera in terms of total number of sequences (*Arsenophonus*, Ca. *Midichloria, Rickettsia, Wolbachia* and *Spiroplasma*), the number of samples that harbored them, beyond 10% of the total number of sequences contained in each sample. For *Arsenophonus*, while this genus corresponded to 30% of the total number of sequences in the dataset, we detected sequences of this genus in only 16.7% of samples. These 16.7% of samples contained 96.5% of the total number of sequences in the dataset for this genus. We found the endosymbiont Ca. *Midichloria*, in 53% of nymphs which contained 90% of the total number of Ca. *Midichloria* sequences. Concerning *Rickettsia*, only 7.8% of samples contained 97% of the total number of sequences. Similarly, 92.1% of sequences belonging to *Wolbachia*, were detected in 12.4% of the samples. Finally, 8.1% of samples harboring *Spiroplasma* contained 67% of the total number of sequences corresponding to this genus in the dataset.

### • Temporal dynamics of the Ixodes ricinus microbiota

#### - Principal component analysis performed on the entire dataset

The principal component analysis was performed to determine the temporal variation of the tick microbiota, based on the relative abundances of microbes (**Figure 2**). The first two principal components PC1 and PC2 explained 24.04% and 11.16% of the total variance, respectively (**Figure 2A**). All samples were clustered into three groups. The first one was projected in the lower right quarter and was opposed to the rest of the analyzed samples, mainly according to the first axis. This small “outsider” cluster was composed of nymphs collected during different months and that seems to be randomly projected regarding the month variable. The remaining samples were divided into two clusters which were represented by ticks collected in March/April and May/June/July/August/September respectively. Note that these two clusters partially overlap.

**Figure 2:**
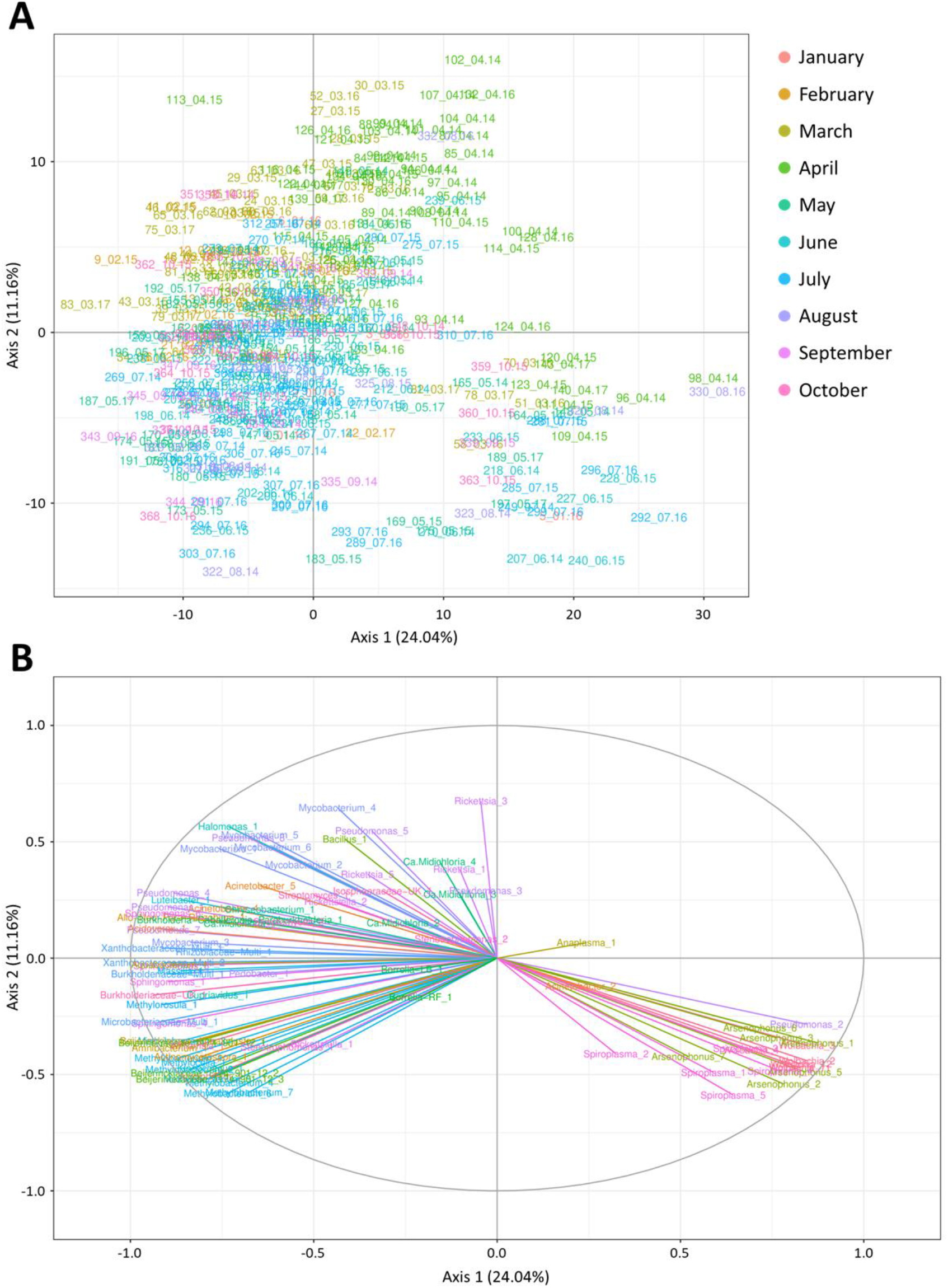
Principal component analysis performed on the entire dataset, presented according to axes 1 (24.04%) and 2 (11.16%). (A) Sample projection of the PCA. Samples are colored according to the month of tick sampling. Plotted samples are named as following: ID_Month.Year. (B) Variable factor map of the PCA. OTUs are colored by genera or by OTUs when belonging to a multi-affiliated (Multi) or unknown (UK) genera.

Four main genera gathering 17 OTUs seem to drive the formation of the “outsider” cluster: *Wolbachia* (number 1, 2, 3, 4, 11, and 12), *Arsenophonus* (number 1, 2, 3, 5, 6 and 7), *Spiroplasma* (number 1, 3, 4 and 5) and *Pseudomonas* (number 2) (**Figure 2B**). The identification of the remaining OTUs driving the two other clusters is more tricky as they are not distinctly separated. In any way, it seems that OTUs belonging to the Beijerinckiaceae (Beijerinckiaceae_alphaI.cluster_1, Beijerinckiaceae_1174-901-12_1, *Methylobacterium*_1-2-3, *Methylorosula*_1, *Methylocella*_1, *Massilia*_1, *Aquabacterium*_1 and *Cupriavidus*_1), Xhantobacteriaceae (Xanthobacteriaceae_Multi_1-2), Burkholderiaceae (Burkholderiaceae-UK_1, Multi_1, *Burkholderia-Caballeronia-Paraburkholderia*_1 and *Acidovorax*_1), Rhizobiaceae (Rhizobiaceae-Multi_1 and *Allo-Neo-Para-Rhizobium*_1) and Microbacteriaceae (Microbacteriaceae-Multi_1, *Amnibacterium*_1) families as well as OTUs *Actinomycetospora*_1, *Williamsia*_1, *Pseudomonas*_1-4-7, *Luteibacter*_4, *Sphingomonas*_1-3-4-6, *Mycobacterium*_1-3 and *Kineococcus*_1 strongly explained the variance along the negative part of the second axis.

#### - Principal component analysis performed on the No Wolbachia/Arsenophonus dataset

This second principal component analysis was performed to determine the temporal variation of the tick microbiota excluding the *Wolbachia* and *Arsenophonus* positive samples as we hypothesized, based on the literature [39,40], that these two genera might not be real members of the tick microbiota (this hypothesis is developed in the discussion part). Here, the first two principal components PC1 and PC2 explained 19.47% and 11.05% of the total variance, respectively (**Figure 3A**). The clustering according to months is still visible with samples clustered into three groups: the cluster 1 mainly composed with samples collected in February/March, the cluster 2 with samples collected in April and the third cluster regrouping mainly samples collected in May/June/July/August/September. Samples collected in October seemed to be distributed in both clusters 1 and 3 (mainly on the left part of the plot, separated from the rest of the community *via* the first axis), while those collected in January do not seem to follow any particular distribution. Ticks collected in April (Cluster 2) are distributed all along the second axis, but only on the right part of the plot, and therefore seemed to be separated from the rest of the community mainly *via* the first axis. By contrast, ticks in the cluster 1, mainly collected in February/March, seemed to be distributed all along the first axis, but mainly on the bottom of the plot, and are consequently separated from the rest of the community mainly through the second axis. For the cluster 3, all samples seemed to be distributed on the top left corner of the plot, distinct from the two other clusters 1 and 2 mainly through the first and second axis, respectively.

**Figure 3:**
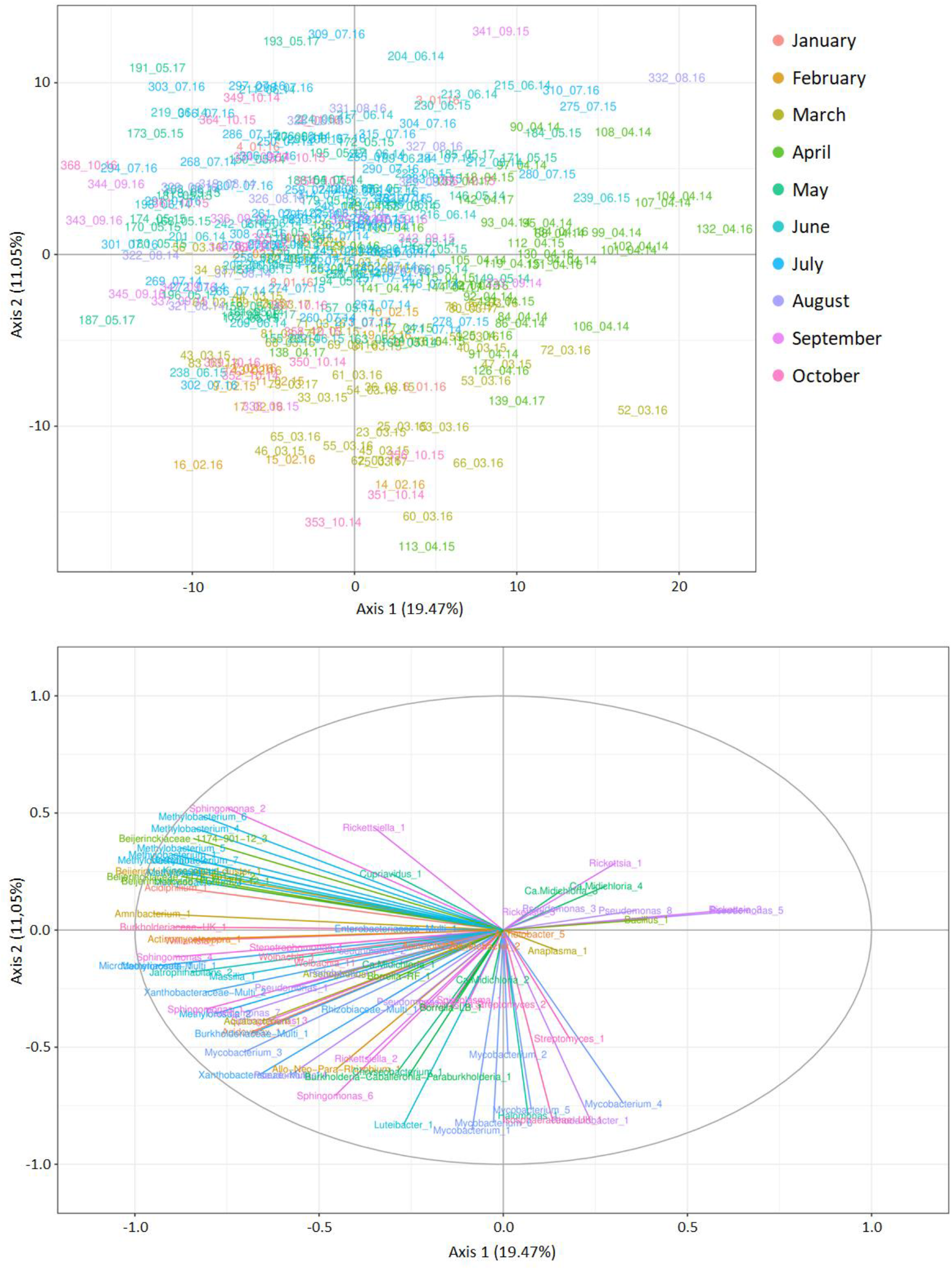
Principal component analysis performed on the dataset excluding *Wolbachia* and *Arsenophonus* positive samples, presented according to axes 1 (20.07%) and 2 (12,08%). (A) Sample projection of the PCA. Samples are colored according to the month of tick sampling. Plotted samples are named as following: ID_Month.Year. (B) Variable factor map of the PCA. OTUs are colored by genera or by OTUs when belonging to a multi-affiliated (Multi) or unknown (UK) genera.

Looking at the variable factor map (**Figure 3B**), we can see that the positive part of the first axis (where are mainly distributed nymphs sampled in April), seemed to be mainly explained by *Rickettsia_3* and *Pseudomonas_5*. Let’s note that their effect is not as strong as those observed on the negative part of the same axis and implicating mainly OTUs belonging to the families Microbacteriaceae (Microbacteriaceae-Multi_1 and *Amibacterium*_1), Beijerinckiaceae (Beijerinckiaceae_1174-901-12_1-2-3, Beijerinckiaceae_Cluster-1, *Methylobacterium*_1-2-3-4-5-6-7, *Methylorosula*_1-2, *Methylocella*_1), Burkholderiaceae (Burkholderiaceae_UK_1), Xhantobacteriaceae (Xhantobacteriaceae-Multi_2), as well as the OTUs *Sphingomonas*_1-4, *Williamsia*_1, *Actinomycetospora*_1, *Jatrophihabitans*_2, *Acidiphilium*-1 and *Kineococcus*_1. The negative part of the second axis (where samples of the Cluster 1 are mainly distributed), seemed to be mainly explained by *Mycobacterium*_1-5-6, Isosphaeraceae-UK_1, *Halomonas*_1, *Rhodanobacter*_1, and *Luteibacter*_1.

### • *Ixodes ricinus* microbiota correlations

Thanks to the network analysis performed on the entire dataset, we observed a total of 219 significant partial correlations between 89 OTUs. Interestingly, 97.8% of these partial correlations were positive (**Figure 4**). Positive partial correlations frequently occurred between OTUs belonging to the same genus or family, as it is the case for OTUs belonging to: *Wolbachia, Arsenophonus, Spiroplasma*, Ca. *Midichloria, Rickettsia, Mycobacterium, Sphingomonas* and Beijerinckiaceae (including Beijerinckiaceae_1174-901-12, Beijerinckiaceae_alphal.cluster, *Methylobacterium, Methylorosula* and *Methylocella*). By contrast, this pattern was not observed between any of the *Pseudomonas* OTUs.

**Figure 4:**
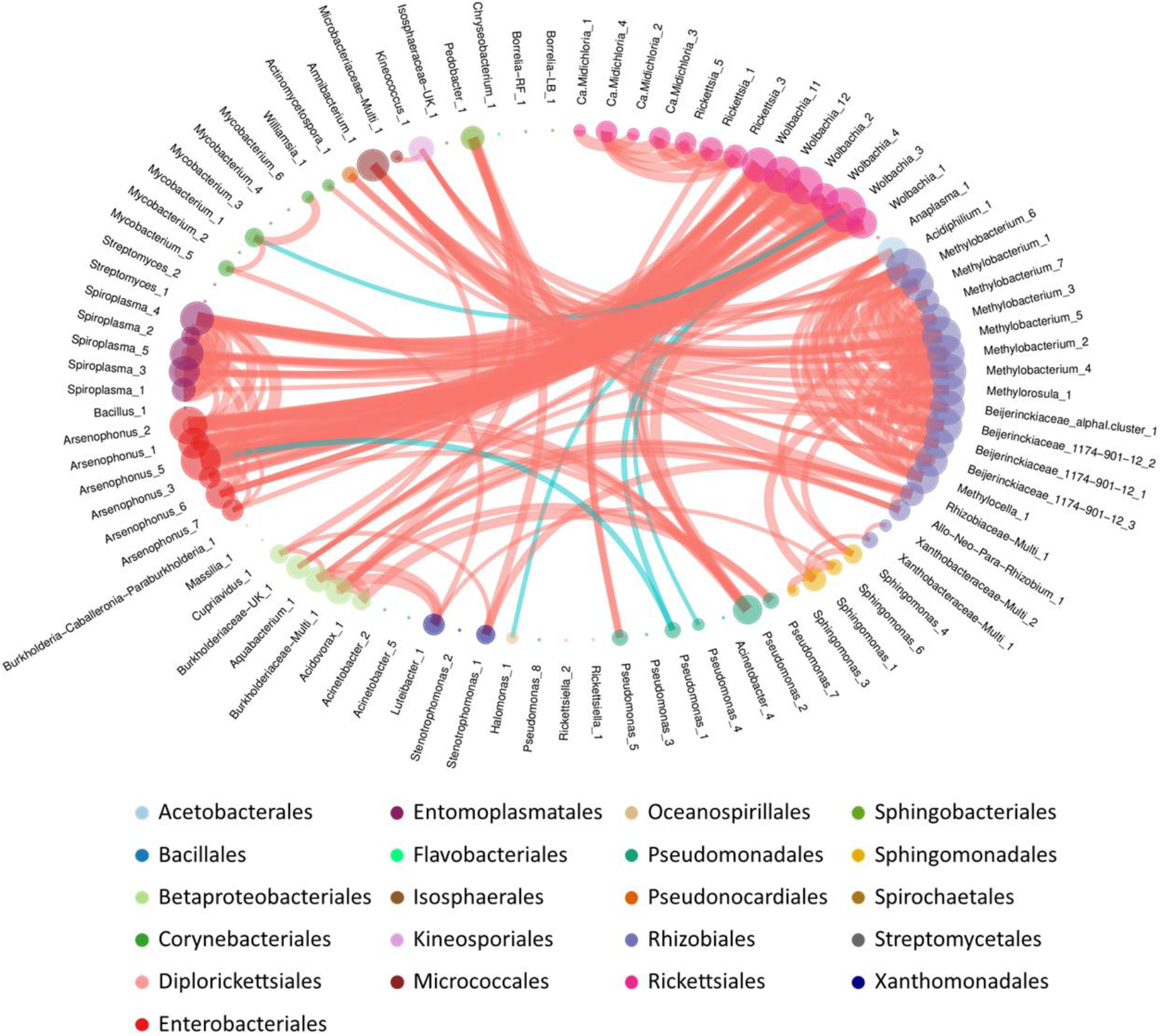
Network analysis. Representation of the significant partial correlations detected between OTUs of the tick microbiota. OTU circles are colored by Taxonomic order. These circles represent nodes of the networks. Their size is proportional to the sum of the incoming edge weights. Thickness of the edge is proportional to the strength of the observed partial correlation. Positive partial correlations are represented by red edges, negative partial correlations are represented by turquoise edges.

In the network, three main clusters were observed. The first one revealed direct connections between several OTUs belonging to *Wolbachia, Arsenophonus* and *Spiroplasma* genera. To these positives partial correlations are added one particular OTU *Pseudomonas*_2, that is linked with several OTUs belonging to *Arsenophonus* and *Wolbachia*. The second main cluster was composed of a large part of OTUs belonging to the Beijerinckiaceae family. Several members of this family also exhibited partial correlations with different OTUs: *Acidiphilium*_1, Burkholderiaceae-UK_1, *Pedobacter* 1, *Kineococcus*_1, *Amnibacterium*_1, *Actinomycetospora*_1, *Williamsia*_1. Finally, a third interesting cluster, smaller than the two others as it implied only 6 OTUs, was characterized by partial correlations between the OTUs Ca. *Midichloria_3-4*, and *Rickettsia*_1-3-5. Let’s note that *Rickettsia*_1-3 are both also linked to *Pseudomonas*_5.

Some negative partial correlations were observed between *Pseudomonas*_1 and both *Arsenophonus*_5 and *Wolbachia*_3. *Wolbachia*_3 was also negatively correlated to *Pseudomonas*_4, *Mycobacterium*_1 and *Halomonas*_1.

### • Links between the presence of tick-borne pathogens and the *Ixodes ricinus* microbiota

#### - Microbiota structure comparison between TBPs positive and negative samples

As expected, the comparison of OTU abundances between each group of TBP positive samples *(Rickettsia, Borrelia* and *Anaplasma*) and negative samples allowed to detect a significantly higher abundance of OTUs belonging to the genus of the tested TBP (**Additional files 1, 2, 3**). Concerning *Rickettsia* positive samples, seven other OTUs were also harboring significantly different abundances compared to negative samples: Ca. *Midichloria_3, Pseudomonas_3-5-8, Bacillus*_1 and Rhizobiaceae-Multi_1 were significantly more abundant in *Rickettsia* positive samples while *Spiroplasma*_1 was significantly less abundant. In Borrelia and Anaplasma positive samples, no other OTUs than those belonging to the corresponding tested genera were significantly over or under represented.

#### - Microbial network comparison between TBPs positive and negatives samples

The network analysis was performed on 5 different datasets. First on samples considered as positive for the presence of *Rickettsia, Borrelia* or *Anaplasma;* then on samples considered as negative for all these genera and finally on all the samples included in the above mentioned datasets, but considering a covariate correcting for the effect of TBPs presence. The determination of Kendall’s τ, integrating the edge appearance rank for all the calculated networks, allowed us to compare them and observe significant differences between all the obtained networks (**Additional file 4**).

In the network analysis performed on positive ticks for *Rickettsia*, we observed 17 significant partial correlations, five were negative **(Figure 5A)**. Several members of the *Rickettsia* genus were positively correlated to OTUs belonging to Ca. *Midichloria* (*Rickettsia*_5/Ca. *Midichlori*a_3) and *Pseudomonas* (*Rickettsia*_1/*Pseudomona*s_3) genera, as already observed in the total dataset network. *Rickettsia*_1, *Pseudomonas*_3 and Ca. *Midichloria*_3 were negatively correlated to *Bacillus*_1. This later OTU was also negatively correlated with 2 OTUs belonging to environmental genera, *Burkholderia*-*Caballeronia*-*Paraburkholderia*_1 and *Chryseobacterium*_1 (that was also positively correlated to Ca. *Midichloria*_3), and positively correlated with *Anaplasma*_1. *Bacillus*_1 therefore appeared as a key member of this network exhibiting several partial correlations with environmental bacteria but particularly with some pathogenic and symbiotic genera. Several positives partial correlations were also observed between OTUs belonging to environmental genera, linking several OTUs within their own genus (*Methylobacterium* and *Mycobacterium*) or between several different genera or families (Beijerickiceae and *Aquabacterium; Stenotrophomonas* and *Acinetobacter* as well as Microbacteriaceae and *Amnibacterium*). We then performed a network analysis considering only ticks positive for *Borrelia* **(Figure 5B).** Only 5 significant partial correlations were observed, including 2 negatives. Negative partial interactions were observed between first *Borrelia*-RF_1 (responsible of Relapsing Fever) and *Borrelia*-LB_1 (responsible of Lyme Borreliosis) and then between *Borrelia*-RF_1 and *Rickettsiella*_2 (**Figure 5B**). A positive partial correlation implying *Spiroplasma*_1 and 2 was also observed and several positive partial correlations implying environmental genera *(Bacillus*_1 and *Massilia*_1 as well as *Mycobacterium_5* and *Luteibacter*_1). In the network analysis performed on ticks only positive for *Anaplasma*, we observed 15 significant partial correlations with 1 identified as negative **(Figure 5C)**. A positive partial correlation was observed between *Rickettsiella*_2 and *Parracoccus*_1. *Spiroplasma*_2 was positively correlated with *Jatrophihabitans*_2 and *Methylobacterium*_3. Other partial correlations concerned environmental bacteria, with members of the Beijerinckiaceae family, correlated with each other or with OTUs belonging to environmental genera corresponding to other families (*Acidiphilium, Allo-Neo-Para-Rhizobium, Amnibacterium* and *Williamsia, Jaotrophihabitans*, Burkholderiaceae). No interactions were observed between *Anaplasma* and the other members of the bacterial community.

**Figure 5:**
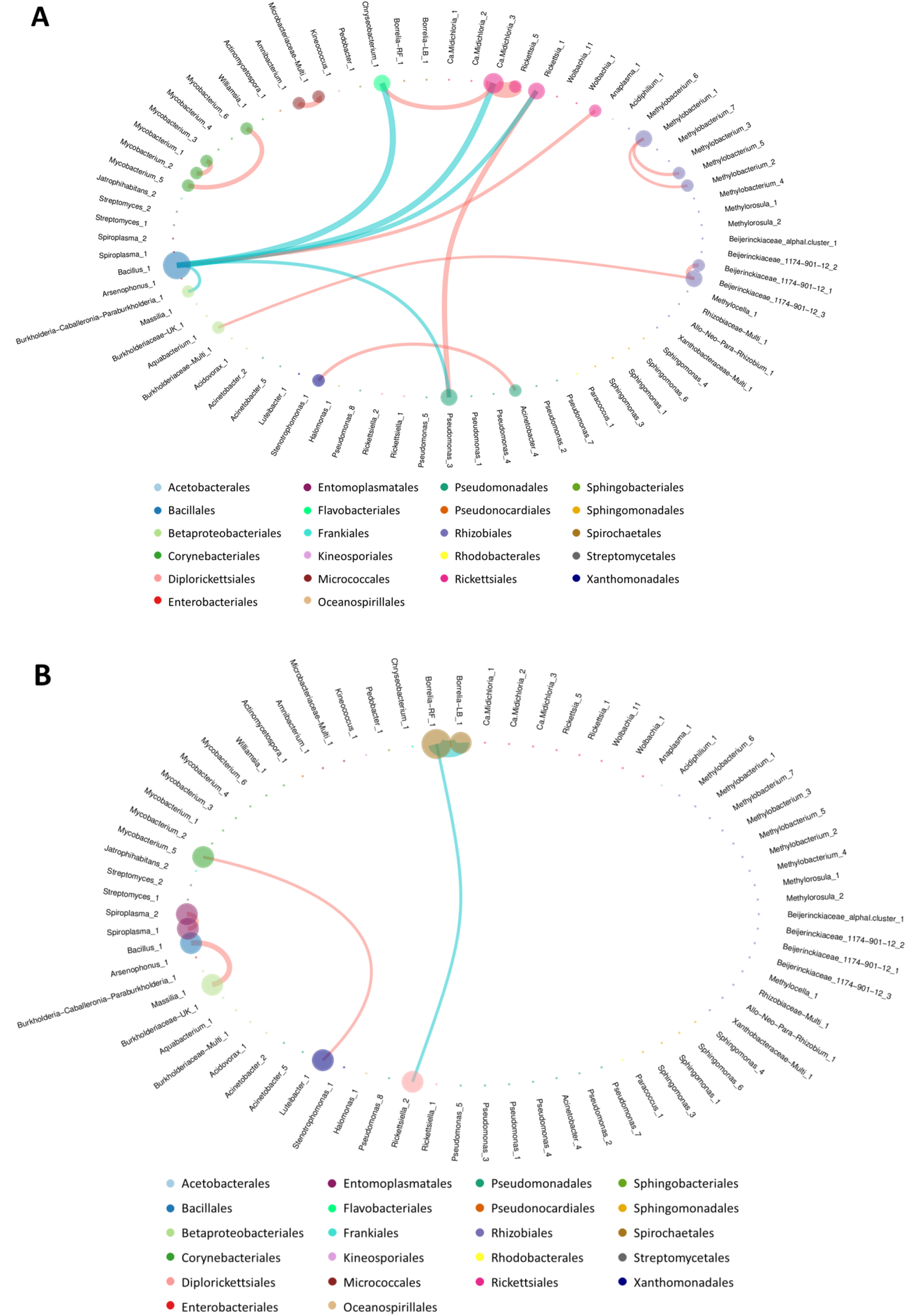
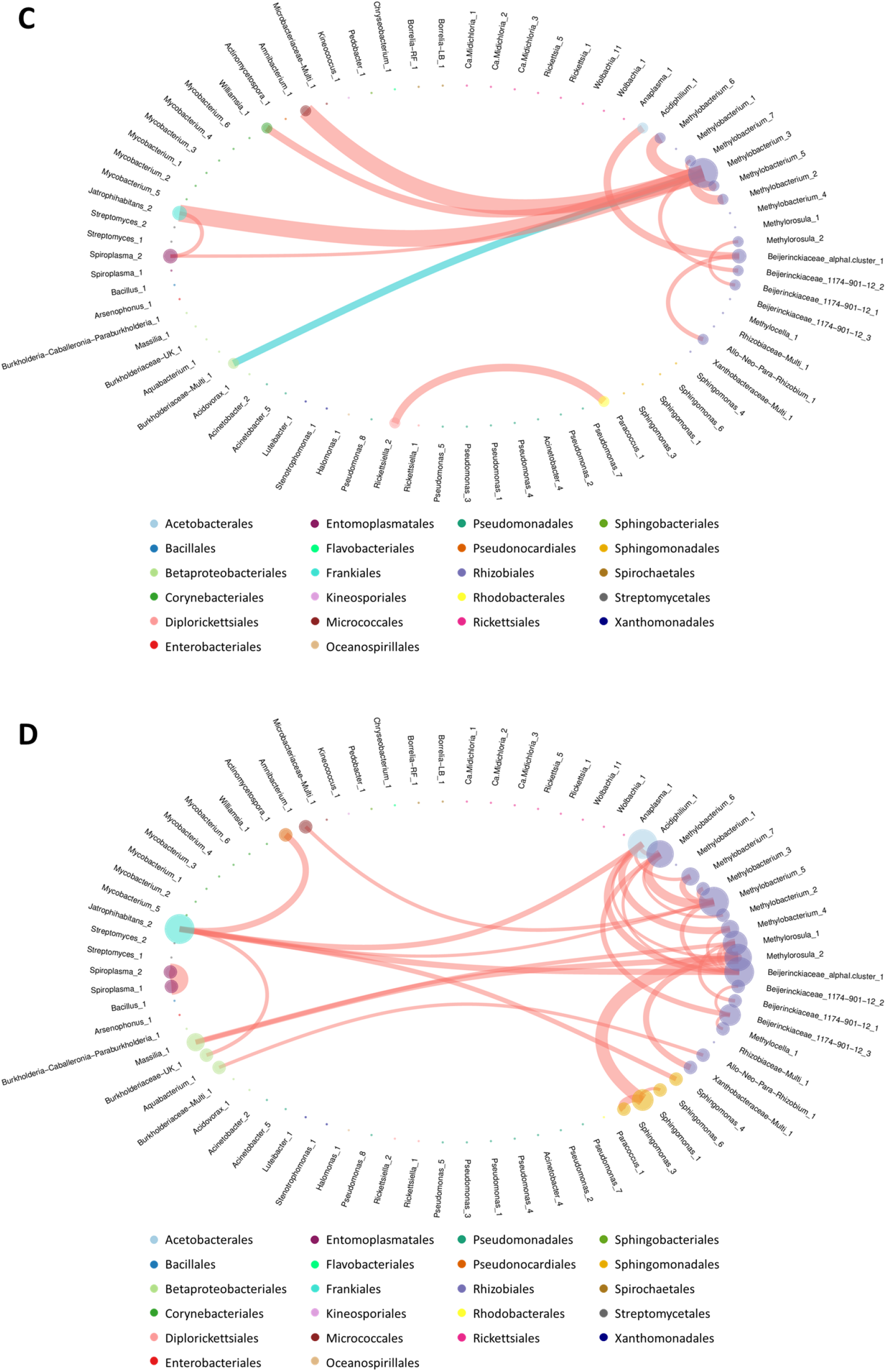
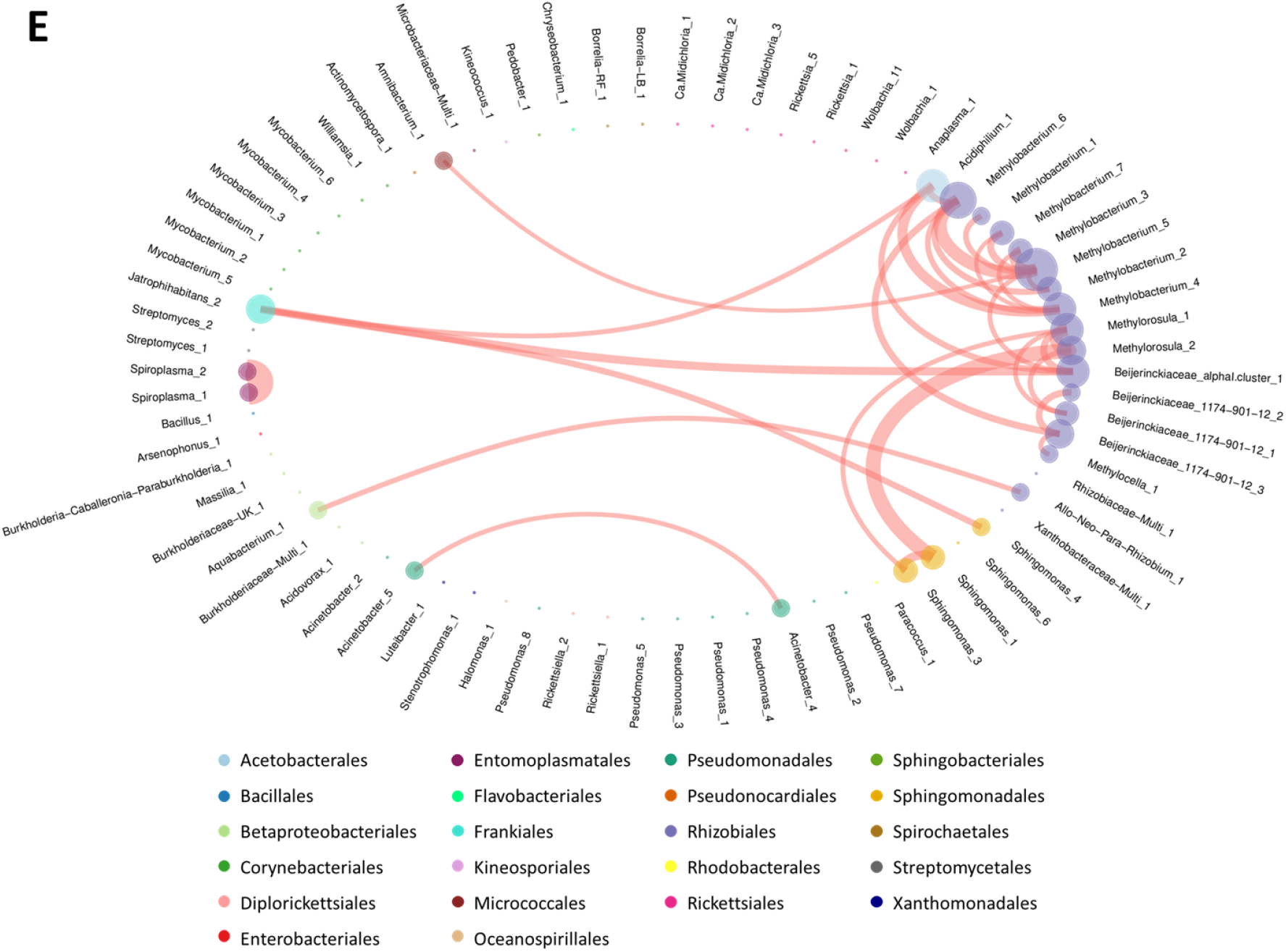
Network analysis. Representation of the significant partial correlations detected between OTUs of the tick microbiota (A) considering only samples positive for Rickettsia, (B) considering only samples positive for Borrelia, (C) considering only samples positive for Anaplasma, (D) considering only negative samples and (E) considering positive and negative samples as well as a covariate accounting for the presence of TBPs. OTU circles are colored by Taxonomic order. These circles represent nodes of the networks. Their size is proportional to the sum of the incoming edge weights. Thickness of the edge is proportional to the strength of the observed partial correlation. Positive partial correlations are represented by red edges, negative partial correlations are represented by turquoise edges.

The negative network **(Figure 5D),** comprising only free pathogen samples, was only composed of positive partial correlations. In this network, a strong partial correlation was observed between *Spiroplasma*_1 and 2. Most of the other correlations were observed between members of the Rhizobiales order, mainly between each other, but also with OTUs belonging to other orders, such as: *Sphingomonas*_1-3, *Acidiphilium*_1, *Jaotrophihabitans*_2, *Aquabacterium*_1, *Massilia*_1, Burkholderiaceae-UK_1, *Amnibacterium*_1. Although it contains less partial correlations, the total network corrected for TBPs effects **(Figure 5E)** is very similar to the negative network. Indeed, partial correlations observed between *Spiroplasma*_1 and 2, as well as most partial correlations involving Rhizobiales order between each other and with other OTUs are also observed in this network.

## Discussion

### • *Ixodes ricinus* microbiota diversity and composition

Investigating the microbiota of the 371 *I. ricinus* nymphs, we identified 109 genera, that is in the range of what it has been previously observed in this tick species [6,12]. Similarly, the mean Shannon diversity index (=2.1) is in the range of what it has been previously observed in the literature for *Ixodes* ticks [8,20,41,42]. However, these values are commonly known to fluctuate, mainly according to the tick stages, species or localization, and are therefore difficult to compare [5,8,10,12,16,42–46]. Furthermore, the fact that not all these studies used negative controls to identify potential contaminating OTUs and remove them from their datasets calls for caution in attempts to draw conclusions about these differences, particularly when we know that these OTUs can represent more than 50% of sequences detected in tick samples [22]. In our dataset, three quarters of the sequences belonged to five genera: *Arsenophonus*, Ca. *Midichloria, Rickettsia, Wolbachia, Spiroplasma*, all of them corresponding to well known maternally inherited bacteria in arthropods [47]. Their high prevalences in the dataset may suggest they would be widely distributed in a large part of tick samples. It was clearly the case for Ca. *Midichloria*, for which most of the detected sequences were found in half of tick samples. This result is completely in concordance with previous observations estimating that almost 100% of *I. ricinus* females and larvae, and almost 50% of nymphs and males were infected by this endosymbiont [48–51]. Concerning the four other genera (*Rickettsia, Spiroplasma, Wolbachia* and *Arsenophonus*), detected sequences were found in less than 17% of samples (7.8%, 8.1%, 12.4% and 16.7% respectively). Focusing on *Rickettsia* sequences, they would correspond to *R. helvetica*, previously detected in these samples [26] and exhibiting a high bacterial load in ticks [52,53]. Concerning *Arsenophonus* and *Wolbachia*, while the high number of sequences detected in the dataset would suggest that they constitute dominant members of the *I. ricinus* microbial communities, their detection might rather be due to the presence of the parasitoid wasp, *Ixodiphagus hookeri* in ticks [39,40]. Finally, among the most detected genera in the *I. ricinus* microbial communities, *Spiroplasma* is a maternally inherited symbiont particularly well known in arthropods, including ticks [54]. With most of their sequences detected in only 8% of tick samples, this bacterial genus would probably represent secondary symbionts [54], which are not essential to their host survival but might confer conditional adaptive advantages to their host, such as protecting their host against pathogens and natural enemies [55,56]. We will see further in this discussion that this is probably the case for *I. ricinus* ticks potentially infected with the parasitoid wasp *Ixodiphagus hookeri*. Among the remaining genera, several are known to circulate in *I. ricinus* ticks, such as *Rickettsiella* (a maternally inherited bacteria in arthropods), *Borrelia* (Lyme Borreliosis and Relapsing Fever) and *Anaplasma*. Some others such as *Methylobacterium, Mycobacterium, Pseudomonas, Stenotrophomonas, Williamsia, Bacillus, Chryseobacterium, Acinetobacter, Allorhizobium-Neorhizobium-Pararhizobium-Rhizobium*, as well as multi-affiliated OTUs belonging to Rhizobiaceae and Microbacteriaceae, previously detected in *Ixodes ricinus* ticks [6,12,16,18,19], were identified as bacteria present in the environment [57–62]. While very few of the above mentioned references reported performing negative controls while studying tick microbiota [12,19], it is important to keep in mind that some of these genera could also correspond to contaminants that may arise during extraction or amplification steps [22,30,63-65]. Here, the dataset underwent a thorough contaminant filtering, based on negative control composition [22], decreasing in this way the risk that these OTUs correspond to contaminants. Nonetheless, questioning remains about these environmental taxa. Do they belong to the internal tick microbiota or to the cuticular microbiome? Probably both. Indeed, Binetruy *et al*. [66], demonstrated that the tick surface sterilization using ethanol (as we did in the present study) did not efficiently remove bacterial DNA from the tick cuticle compared to bleach. They also showed that environmental taxa, only detected in ethanol cleaned entire ticks, belonged to the same family than those directly detected on the tick cuticle by swabbing. However, they also found some of these taxa in the gut of ticks and Hernández-Jarguín *et al*. [18], while studying the microbiota in organs of *I. ricinus* ticks previously cleaned with bleach, also detected bacterial genera commonly found in the environment.

### • Temporal variability of Ixodes ricinus microbiota

We first performed a principal component analysis on the entire dataset to characterize the temporal dynamics of the *I. ricinus* microbiota. Our results provided evidence that the *I. ricinus* microbial communities exhibit marked temporal variations with two main clusters, a first one defined by microbial communities identified in March-April and another one with microbial communities identified from May to September. Interestingly, this temporal pattern was annually recurrent suggesting predictable temporal differences in the *I. ricinus* microbial community structure. A third cluster was also identified and corresponded to nymphs collected through the three years. This “outsider” cluster was mainly composed of members belonging to the genera *Wolbachia, Arsenophonus* and *Spiroplasma*. As previously mentioned, the presence of species belonging to *Wolbachia* and *Arsenophonus* in *I. ricinus* ticks is probably due to the presence of the parasitoid wasps, *Ixodiphagus hookeri* [39,40], that is strictly associated with tick for its development. In their study, Plantard *et al*. [39] reported that *I. hookeri* was infested at almost 100% prevalence by *Wolbachia pipientis*. They also showed that all unfed nymphs parasitized by *I. hookeri* also harboured *Wolbachia*, while non infected ticks were *Wolbachia-free* (with only one exception). *Wolbachia* was vertically transmitted from the female *I. hookeri* to its eggs and the subsequent generation harboured *Wolbachia*. Similarly, Bohacsova *et al*. [40] detected the symbiont *Arsenophonus nasoniae* only in nymphs infected by the wasp *I. hookeri*, whose population was harboring at almost 30% prevalence with *A. nasoniae*. In this context, through the study of tick microbial communities, we suggest that identifying the temporal dynamics of both *Wolbachia* and *Arsenophonus* could serve as a proxy to characterize both the infection rate and temporal dynamics of *I. hookeri* in *I. ricinus*, even if it would probably be easier and more efficient to use primers or probes specifically matching with the parasitoid DNA. While the presence of *Wolbachia* and *Arsenophonus* was thus probably linked to the presence of *I. hookeri* in ticks, we hypothesized that their presence in our dataset could alter the characterization of the temporal dynamics of the *I. ricinus* microbial communities. We therefore chose to perform another PCA analysis after removing all samples harboring one or both genera *Wolbachia* and *Arsenophonus*. As expected, the new PCA performed on the “reduced” dataset allowed to improve the separation between clusters. We finally distinguished three main clusters. Whatever the year of sampling, this analysis mainly distinguished ticks collected in February/March (cluster 1), from those sampled in April (cluster 2) or those sampled from May to September (Cluster 3), with ticks sampled in October distributed with both cluster 1 and 3. Interestingly, this pattern was still likely to be annually redundant, suggesting that factors driving tick microbial communities would be probably the same during the three consecutive years. Interestingly, we observed that the OTUs that better explained the dynamics of the tick microbial community structure were not tick symbionts nor pathogens but corresponded to genera or families characteristics of the environment (i.e. *Sphingomonas, Williamsia, Actinomycetospora, Jatrophihabitans, Acidiphilium, Kineococcus, Mycobacterium, Halomonas*, Microbacteriaceae, Beijerinckiaceae, Xhantobacteriaceae, Burkholderiaceae, Rhodanobacteraceae and Isosphareaceae). Considering that these OTUs would belong to the cuticular microbiome (as discussed above), the observed fluctuations could therefore correspond to variations of the environmental microbiota [67]. Nonetheless, these OTUs could also correspond to bacteria acquired by ticks from the environment, during the water uptake mechanism for example, or from other orifices of the tick such as spiracles and anal port. This was already suggested by Narasimhan and Fikrig [68] while they observed that ticks hatched in a sterile environment harbored a significantly different gut microbiota than those hatched in “normal” conditions [69]. In such a case, temporal variations observed in our analysis would also be due to differences of environment microbiota composition. Otherwise, abiotic factors such as temperature, that is likely to follow a yearly redundant pattern, have been demonstrated to influence the microbiota diversity of *I. scapularis* ticks [46]. Furthermore, blood meal and host identity were also shown to influence the tick microbiota diversity [11,41,69,70]. Finally, considering the life cycle of ticks, one other potential hypothesis explaining this pattern is that tick microbiota structure and composition could be related to the feeding status of ticks. While quantifying the lipidic content in *Ixodes ricinus* ticks monthly sampled over several years, Randolph *et al*. [71] and Abdullah *et al*. [72] managed to discriminate tick feeding cohorts questing at different periods of the year. Both of them noticed that lipidic reserves were globally higher in ticks collected at the end of the year, and lower for those sampled from the end of spring and through summer. According to Abdullah *et al*. [72], ticks sampled at the end of the year (high lipidic reserve), would correspond to those managing to perform their blood meal during the last spring and to emerge in Fall. At the beginning of the next year (between the end of winter and the beginning of the next spring), sampled ticks would correspond to a mix of tick populations: 1) those just emerging, exhibiting high lipidic reserve and 2) those that already emerged in fall, still exhibiting pretty high lipidic level. Ticks still questing at the end of spring or during summer would correspond to those that failed to find a host and almost exhausted their lipidic reserves. Considering this, one could hypothesize that ticks constituting the cluster 3 (mainly ticks from May to September) could correspond to those with very few energetic reserves, while the cluster 1 (ticks from February/March) would correspond to those presenting higher energetic reserves. Ticks sampled in October, distributed among the clusters 1 and 3, could correspond to a mix of two populations: 1) those with high lipidic reserves (that fed during last spring and just emerged) and 2) those with almost exhausted energetic reserve (that fed the previous year), as observe for nymphs sampled in October 2015 in Abdullah *et al*. [72]. However, this hypothesis does not allow to elucidate the clustering of ticks sampled in April suggesting that other factors (related to the life cycle of ticks or their environment), could be at the origin of this pattern. Further studies will be necessary to elucidate this point and confirm or not this hypothesis.

### • *Ixodes ricinus* microbial community interactions

Identifying and understanding tick-borne microbe interactions represents an essential prerequisite to develop new control strategies of ticks and tick-borne diseases using the tick microbiome. Using partial correlation networks, we assessed which tick bacterial OTUs were correlated to identify possible associations between tick microbial community members.

First, considering the entire dataset, the network analysis revealed that more than 97% of interactions between members of the *I. ricinus* microbial community were positive. Taxa with positive associations have generally been interpreted as functional guilds of organisms performing similar or complementary functions [73,74] or feature interactions shaped by interspecies cross-feeding [75], although sometimes they may, to a large extent, reflect similar preferred conditions [76]. Similarly, negative associations might reflect interactions including competition and niche partitioning. The vast majority of positive correlations observed could therefore suggest that tick microbial communities might favor mutualism and perform similar or complementary functions. But let’s keep in mind that correlations observed during this study were obtained by examining entire individuals, and may impair the observation of associations occurring at a finer scale (e.g. organs) [27].

Due to the low infection rate observed in studied nymphs [26], the low representation of positive ticks for pathogens in our dataset might distort the analyses and conceal important information as crucial interactions between TBPs and members of the tick microbiota To overcome this bias, we decided to perform supplemental analysis, comparing TBPs positive and negative samples. Furthermore, in order to get rid of effects that could hamper or skew the observation of TBPs impact, samples that appeared to be parasitized with *I. hookerii* (i.e. presenting a numerous number of sequences belonging to *Arsenophonus* and/or *Wolbachia*) were excluded from this analysis, and a covariate correcting for the effect of sampling season was added, compared to the analysis performed on the entire dataset. The obtained negative network was mainly composed of partial correlations linking environmental OTUs and very few correlations remain compared to the network obtained with the entire dataset. These results suggest that most correlations previously observed in the total network are linked to the presence of TBPs, the response to parasitoid or the seasonal effect. This strengthens once again the importance of these variables in the tick microbial community structure and underlines the adaptability of the tick microbiota to variable conditions. Comparing negative samples with those infected by pathogens allowed us to demonstrate that with TBPs, the structure and correlations of tick microbial communities were considerably modified. Indeed, in ticks infected by *Rickettsia*, proportions of several OTUs (i.e. *Spiroplasma* symbiont and environmental OTUs) were significantly affected. Moreover, more negative correlations were detected in TBPs samples, suggesting much more competition between members of the tick microbial communities in presence of TBPs. In addition, in the “free pathogen” network, we observed a lot of correlations implying environmental OTUs (i.e. belonging to Rhizobiales and Betaproteobacteriales orders as well *Sphingomonas* genera and *Jatrophihabitans*_2), whereas these correlations decreased or disappeared in infected ticks, suggesting that the presence of TBPs would affect these microbial interactions. Two main hypotheses emerged from these results: the tick microbiota might be initially disturbed and thus favor the infection by pathogens or the presence of pathogens might impact the structure of the tick microbiota. Because the structure and interactions of tick microbial communities are completely different according to the pathogen considered, we could first suggest that the presence of pathogens in ticks would be likely to affect the other members of the tick microbial communities. However, the microbiota disturbance might facilitate the installation of pathogens in ticks. As an example, the presence of *Anaplasma phagocytophilum* in *I. scapularis* would modify the gut microbiota whose modification would disrupt the integrity of the gut membrane and help the pathogen entry [77]. Based on our data, we are currently not able to conclude and experimental approaches will have to be performed in the future.

#### - Interactions between non-pathogenic OTUs

We first observed strong correlations between *Wolbachia, Arsenophonus* and *Spiroplasma* OTUs. As previously mentioned, detecting *Wolbachia* and *Arsenophonus* in *I. ricinus* probably betrayed the presence of the parasitoid *I. hookeri* in these ticks [39,40]. Interestingly, *Spiroplasma*, identified as arthropod symbionts including ticks [47,54,78–81], were highly linked to the dynamics of these two genera. Identified as a “male killer” in many other arthropods [82–87], our results only on nymphs do not allow here to conclude about this potential role even if no evidence of sex-ratio distortion has been observed in the *Rhipicephalus decoloratus* tick population infected with *Spiroplasma ixodetis* [54]. Assuming that the detection of both *Wolbachia* and *Arsenophonus* is probably due to the infection of ticks by *I. hookeri*, we hypothesize that *Spiroplasma* might be upregulated by the presence of the parasitoid in ticks and might act as a defensive response mechanism against *I. hookeri*, as previously mentioned in *Drosophila melanogaster* infested by two species of parasitoid wasps [88]. Nevertheless, we cannot rule out the hypothesis that, while identified in a large variety of tick species [47,54], *Spiroplasma* could correspond to a symbiont of *I. hookeri*. Let’s note that contrary to *Spiroplasma*, several *Pseudomonas* OTUs were either negatively or positively correlated to *Wolbachia* and *Arsenophonus*. In our dataset, very few OTUs belonging to the same genus displayed contrasting patterns of correlation, but *Pseudomonas* is an exceptionally versatile genus including plant pathogens, human or animal opportunistic pathogens and saprophytic bacteria found in water and soil exhibiting great adaptation to their environment [89]. Some members of this genus have notably been reported to have an influence on arthropod survival. For example, *P. fluorescens* strains were reported to confer better survival to *Varroa destructor* mite and *Galleria mellonella* waxworm in presence of a fungal pathogen of arthropods *Beauveria bassiana* [90,91]. On the opposite, *P. entomophila* was reported to be entomopathogenic for *Drosophila* [92,93]. Such opposite behavior in arthropods could explain the contrasting correlations observed with *Pseudomonas* OTUs, and suggest that, as *Spiroplasma* OTUs, some of them might be implied in defense mechanisms against *I. hookeri*i.

Another group of OTUs, including *Methylobacterium*, OTUs of the Beijerinckiaceae family, *Acidiphilium*_1, Burkholderiaceae-UK_1, *Pedobacter_1, Kineococcus*_1, Microbacteriaceae-Multi_1, *Amnibacterium*_1, *Actinomycetospora*_1, *Williamsia*_1, exhibited a lot of positive interactions. All these OTUs belonged to genera and families of microorganisms commonly found in the environment [60,94–99], and known to potentially infect ticks. The high connectivity displayed by these OTUs might reflect similar functions between OTUs adapted to contrasting environmental conditions (functional redundancy). However, such a scenario would have probably reflected much more negative correlations between these OTUs. On the contrary, these findings might indicate reduced functional capabilities of each OTU and thus a complementary functional role of these taxa in ticks. All these hypotheses will have to be investigated in future works studying the functional ecology of tick microbes.

#### - Interactions between tick-borne microbiota and pathogens

We interestingly observed correlations between Ca. *Midichloria* and *Rickettsia* OTUs, strong enough to be observed not only in the *Rickettsia* network, but also in the total dataset networks. While our sequencing approach does not allow us to identify *Rickettsia* at the species level, they would correspond to pathogenic agents as *Rickettsia helvetica* was specifically detected in these same samples [26]. In addition, Ca. *Midichloria* abundance was significantly higher in *Rickettsia* positive samples compared to the negative ones. Considering the strong prevalence of Ca. *Midichloria* in *I. ricinus* ticks, the positive relationship between this OTU and *Rickettsia* might suggest a facilitating role of Ca. *Midichloria* in the *I. ricinus* colonisation by *Rickettsia*. This hypothesis is in accordance with Budachetri *et al*.’s results [28] which observed positive correlation between Ca. *Midichloria mitochondrii* load and *Rickettsia parkeri* presence in the tick *Amblyomma maculatum*. The fact that *Rickettsia* were positively correlated with a maternally inherited bacteria is particularly surprising, especially because members of this genus have been frequently reported to be involved in antagonist relationships with symbiotic or pathogenic genera [9,10,19,100–103]. This potential complementarity between these two genera will have to deepen in the future by characterizing the bacterial transcriptome or metabolome of ticks infected with these bacteria alone or in association. The *Borrelia* Relapsing fever OTU (*Borrelia* RF) was interestingly negatively correlated with *Rickettsiella* and *Borrelia* Lyme Borreliosis OTUs (*Borrelia* LB). These negative correlations between both *Borrelia* groups (Relapsing Fever *vs* Lyme Borreliosis) using the 16S rDNA are completely in accordance with our previous results obtained from the same tick samples detecting specifically tick-borne pathogens using a high-throughput microfluidic real-time PCR with specific primers and probes [26]. Indeed, even though this association was not significantly underrepresented compared to a random distribution, our previous results highlighted higher prevalences of *Borrelia burgdorferi s.l*. when *B. miyamotoi* prevalences were low and vice versa. Competition for the same niche might explain the negative correlations observed between these two groups of *Borrelia*. However, these findings contrast with those presented in Aivelo *et al*. [19] as positive correlations were detected between the relapsing fever spirochete *B. miyamotoi* with *Rickettsiella* and two *Borrelia* species belonging to the group *Borrelia* Lyme borreliosis (*B. garinii, B. afzelii*). Because clinical co-infections with several TBPs are commonly reported [104–106] and are known to affect both disease symptoms and severity [107,108], it will be crucial to determine conditions in which *Borrelia* RF and *Borrelia* LB would be in competition or on the contrary could collaborate and thus coinfect ticks.

We also observed several partial correlations between pathogens and OTUs usually known to be “environmental” bacteria. It was notably the case regarding a *Pseudomonas* OTU positively correlated with *Rickettsia*. In addition, this OTU and two others belonging also to this genus showed higher abundances in *Rickettsia* positive samples. While several *Pseudomonas* OTUs were previously identified as contaminants and removed from the analysed dataset [22], those remaining are implied in several interactions with different members of the tick microbial community, such as *Wolbachia* and *Arsenophonus*, as already discussed, but also to TBPs, demonstrating the versatility of members of this genus and their importance within the tick microbiota structure. *Bacillus* also appears as a key member of the *Ixodes ricinus* microbial community linked to the presence of TBPs. While Adegoke *et al*. [109] observed higher abundances of this bacterial genera with the presence of the parasite *Theileria* in the tick *Rhipicephalus microplus*, our findings demonstrate a positive correlation between *Bacillus* and *Anaplasma* and a negative one with *Rickettsia*. These latest would suggest potential competition between these bacteria and might be a first information to develop a future tool to control tick infections by *Rickettsia*. Furthermore, because the dynamics of environmental bacteria found in ticks varies through the year probably with the contrasting environmental conditions, the vegetation and the tick hosts, their positive or negative interactions with pathogens might suggest that their presence or absence would be an important factor to take into account to better understand the temporal dynamics of TBPs. Finally, all these correlations implying “environmental” OTUs suggest that their detection in tick microbiota is probably not only the result of accidental ingestion, and is more likely to reflect their true adaptation within the tick microbial community.

## Conclusions

Here we reported the identification of the *Ixodes ricinus* microbiota from nymphs collected monthly over several consecutive years. Results allowed to show that 1) the *Ixodes ricinus* microbiota is not stable over time but follow a recurrent temporal pattern, mainly explained by the dynamics of environmental taxa; 2) the presence of TBPs is likely to disturb the tick microbial community structure and thus tick microbe interactions; 3) some specific symbionts and “environmental” bacteria might play a key role on the presence and the dynamics of *I. ricinus-borne* pathogens and in the defense against parasitoid species. While microbe correlations identified from this ecosystemic study will have to be confirmed in the next future using experimental approaches, our new findings suggest that all members of the tick microbiome, including environmental taxa, would play a key role in both the *Ixodes ricinus* fitness and the presence of tick-borne pathogens and should be considered as a promising tool for the development of new controls strategies against tick and tick-borne diseases.

## List of abbreviations

*I. ricinus*: *Ixodes ricinus*
PCA: Principal Component Analysis
OTU: Operational Taxonomic Unit
TBP: Tick-Borne Pathogens

## Declarations

## Availability of data and material

The datasets used for this study can be found in the European Nucleotide Archive. Project accession number: PRJEB36903 (ERP120162) – Sample accession numbers: ERS4353953-ERS4354625 – Run accession numbers: ERR3956669-ERR3957340.

## Competing interests

Author Cédric Midoux was employed by the company Irstea. The remaining authors declare that the research was conducted in the absence of any commercial or financial relationships that could be construed as a potential conflict of interest.

## Funding

This work was supported by the métaprogramme “Metaomics and microbial ecosystems” (MEM) and the métaprogramme “Adaptation of Agriculture and Forests to Climate Change” (ACCAF) granted by the French National Institute for Agricultural Research (France). The salary of Emilie Lejal, the PhD student working on this project was funded by the Ile-de-France region. Julie Aubert, Julien Chiquet and Stéphane Robin were partially supported by the French ANR grant ANR-17-CE32-0011 Next Generation Biomonitoring (NGB).

## Acknowledgements

This work was supported by the métaprogramme “Metaomics and microbial ecosystems” (MEM) and the Métaprogramme “Adaptation of Agriculture and Forests to Climate Change” (ACCAF) granted by the French National Institute for Agriculture, Alimentation and Environment Research (France). The authors would like to thank Sabine Delannoy and the IdentyPath genomic platform that allowed us to perform part of the experiments in their laboratory. We also thank Maxime Galan for providing index sequences and stimulating discussions. We are grateful to the INRAE MIGALE bioinformatics facility (MIGALE, INRAE, 2020. Migale bioinformatics Facility, doi: 10.15454/1.5572390655343293E12) for providing help and computing and storage resources. Finally, the authors would like to thank Olivier Plantard for enriching discussions particularly on the links between tick microbiota and energetic reserves.

## Additional files

**Additional file 1: Microbiota structure comparison between *Rickettsia* positive and negative samples.** Differential analysis, comparing OTUs abundances between *Rickettsia* positive and negative samples, considering a 95% confidence interval. Pvalues are corrected according to Bonferroni adjustment.

**Additional file 2: Microbiota structure comparison between *Borrelia* positive and negative samples.** Differential analysis, comparing OTUs abundances between *Borrelia* positive and negative samples, considering a 95% confidence interval. Pvalues are corrected according to Bonferroni adjustment.

**Additional file 3: Microbiota structure comparison between *Anaplasma* positive and negative samples.** Differential analysis, comparing OTUs abundances between *Anaplasma* positive and negative samples, considering a 95% confidence interval. Pvalues are corrected according to Bonferroni adjustment.

**Additional file 4: TBPs positive and negative network comparison.** The different families of network generated in this study (negative, *Rickettsia, Borrelia, Anaplasma* and total corrected for TBPs effect) were compared by calculating a weighted version of the Kendall’s τ, integrating the edge appearance rank within families of networks (negative, *Rickettsia, Borrelia, Anaplasma* and total corrected for TBPs effects) as well as associated Pvalues.

## Notes

### Competing Interest Statement

Author Cedric Midoux was employed by the company Irstea. The remaining authors declare that the research was conducted in the absence of any commercial or financial relationships that could be construed as a potential conflict of interest.

## References

1. Duron O, Noël V, McCoy KD, Bonazzi M, Sidi-Boumedine K, Morel O, et al. The Recent Evolution of a Maternally-Inherited Endosymbiont of Ticks Led to the Emergence of the Q Fever Pathogen, Coxiella burnetii. PLOS Pathogens. 2015;11:e1004892.

2. Duron O, Morel O, Noël V, Buysse M, Binetruy F, Lancelot R, et al. Tick-Bacteria Mutualism Depends on B Vitamin Synthesis Pathways. Current Biology. 2018;28:1896–1902.e5.

3. Vayssier-Taussat M, Kazimirova M, Hubalek Z, Hornok S, Farkas R, Cosson J-F, et al. Emerging horizons for tick-borne pathogens: from the ‘one pathogen–one disease’ vision to the pathobiome paradigm. Future Microbiology. 2015;10:2033–43.

4. Bonnet SI, Binetruy F, Hernández-Jarguín AM, Duron O. The Tick Microbiome: Why Non-pathogenic Microorganisms Matter in Tick Biology and Pathogen Transmission. Front Cell Infect Microbiol [Internet]. 2017 [cited 2019 Aug 30];7. Available from: https://www.frontiersin.org/articles/10.3389/fcimb.2017.00236/full

5. Lalzar I, Harrus S, Mumcuoglu KY, Gottlieb Y. Composition and Seasonal Variation of Rhipicephalus turanicus and Rhipicephalus sanguineus Bacterial Communities. Appl Environ Microbiol. 2012;78:4110–6.

6. Nakao R, Abe T, Nijhof AM, Yamamoto S, Jongejan F, Ikemura T, et al. A novel approach, based on BLSOMs (Batch Learning Self-Organizing Maps), to the microbiome analysis of ticks. The ISME Journal. 2013;7:1003–15.

7. Zhang X-C, Yang Z-N, Lu B, Ma X-F, Zhang C-X, Xu H-J. The composition and transmission of microbiome in hard tick, Ixodes persulcatus, during blood meal. Ticks and Tick-borne Diseases. 2014;5:864–70.

8. Van Treuren W, Ponnusamy L, Brinkerhoff RJ, Gonzalez A, Parobek CM, Juliano JJ, et al. Variation in the Microbiota of Ixodes Ticks with Regard to Geography, Species, and Sex. Appl Environ Microbiol. 2015;81:6200–9.

9. Gall CA, Reif KE, Scoles GA, Mason KL, Mousel M, Noh SM, et al. The bacterial microbiome of Dermacentor andersoni ticks influences pathogen susceptibility. The ISME Journal. 2016;10:1846–55.

10. René-Martellet M, Minard G, Massot R, Tran Van V, Valiente Moro C, Chabanne L, et al. Bacterial microbiota associated with Rhipicephalus sanguineus (s.l.) ticks from France, Senegal and Arizona. Parasites & Vectors. 2017;10:416.

11. Swei A, Kwan JY. Tick microbiome and pathogen acquisition altered by host blood meal. The ISME Journal. 2017;11:813–6.

12. Estrada-Peña A, Cabezas-Cruz A, Pollet T, Vayssier-Taussat M, Cosson J-F. High Throughput Sequencing and Network Analysis Disentangle the Microbial Communities of Ticks and Hosts Within and Between Ecosystems. Front Cell Infect Microbiol. 2018;8:236.

13. Varela-Stokes AS, Park SH, Stokes JV, Gavron NA, Lee SI, Moraru GM, et al. Tick microbial communities within enriched extracts of Amblyomma maculatum. Ticks and Tick-borne Diseases. 2018;9:798–805.

14. Tokarz R, Tagliafierro T, Sameroff S, Cucura DM, Oleynik A, Che X, et al. Microbiome analysis of Ixodes scapularis ticks from New York and Connecticut. Ticks and Tick-borne Diseases. 2019;10:894–900.

15. Cabezas-Cruz A, Pollet T, Estrada-Peña A, Allain E, Bonnet SI, Moutailler S. Handling the Microbial Complexity Associated to Ticks. Ticks and Tick-Borne Pathogens [Internet]. 2018 [cited 2019 Sep 1]; Available from: https://www.intechopen.com/books/ticks-and-tick-borne-pathogens/handling-the-microbial-complexity-associated-to-ticks

16. Carpi G, Cagnacci F, Wittekindt NE, Zhao F, Qi J, Tomsho LP, et al. Metagenomic Profile of the Bacterial Communities Associated with Ixodes ricinus Ticks. PLOS ONE. 2011;6:e25604.

17. Gofton AW, Oskam CL, Lo N, Beninati T, Wei H, McCarl V, et al. Inhibition of the endosymbiont “Candidatus Midichloria mitochondrii” during 16S rRNA gene profiling reveals potential pathogens in Ixodes ticks from Australia. Parasites & Vectors. 2015;8:345.

18. Hernández-Jarguín A, Díaz-Sánchez S, Villar M, de la Fuente J. Integrated metatranscriptomics and metaproteomics for the characterization of bacterial microbiota in unfed Ixodes ricinus. Ticks and Tick-borne Diseases. 2018;9:1241–51.

19. Aivelo T, Norberg A, Tschirren B. Bacterial microbiota composition of Ixodes ricinus ticks: the role of environmental variation, tick characteristics and microbial interactions. PeerJ. PeerJ Inc.; 2019;7:e8217.

20. Portillo A, Palomar AM, Toro M de, Santibáñez S, Santibáñez P, Oteo JA. Exploring the bacteriome in anthropophilic ticks: To investigate the vectors for diagnosis. PLOS ONE. 2019;14:e0213384.

21. Guizzo MG, Neupane S, Kucera M, Perner J, Frantová H, da Silva Vaz IJ, et al. Poor Unstable Midgut Microbiome of Hard Ticks Contrasts With Abundant and Stable Monospecific Microbiome in Ovaries. Front Cell Infect Microbiol [Internet]. Frontiers; 2020 [cited 2020 Sep 21];10. Available from: https://www.frontiersin.org/articles/10.3389/fcimb.2020.00211/full?report=reader

22. Lejal E, Estrada-Peña A, Marsot M, Cosson J-F, Rué O, Mariadassou M, et al. Taxon Appearance From Extraction and Amplification Steps Demonstrates the Value of Multiple Controls in Tick Microbiota Analysis. Front Microbiol [Internet]. Frontiers; 2020 [cited 2020 Sep 14];11. Available from: https://www.frontiersin.org/articles/10.3389/fmicb.2020.01093/full

23. Gassner F, van Vliet AJH, Burgers SLGE, Jacobs F, Verbaarschot P, Hovius EKE, et al. Geographic and Temporal Variations in Population Dynamics of Ixodes ricinus and Associated Borrelia Infections in The Netherlands. Vector-Borne and Zoonotic Diseases. 2010;11:523–32.

24. Coipan EC, Jahfari S, Fonville M, Maassen C, van der Giessen J, Takken W, et al. Spatiotemporal dynamics of emerging pathogens in questing Ixodes ricinus. Front Cell Infect Microbiol. 2013;3:36.

25. Takken W, van Vliet AJH, Verhulst NO, Jacobs FHH, Gassner F, Hartemink N, et al. Acarological Risk of Borrelia burgdorferi Sensu Lato Infections Across Space and Time in The Netherlands. Vector-Borne and Zoonotic Diseases. 2016;17:99–107.

26. Lejal E, Marsot M, Chalvet-Monfray K, Cosson J-F, Moutailler S, Vayssier-Taussat M, et al. A three-years assessment of Ixodes ricinus-borne pathogens in a French peri-urban forest. Parasites & Vectors. 2019;12:551.

27. Pollet T, Sprong H, Lejal E, Krawczyk AI, Moutailler S, Cosson J-F, et al. The scale affects our view on the identification and distribution of microbial communities in ticks. Parasites Vectors. 2020;13:36.

28. Budachetri K, Kumar D, Crispell G, Beck C, Dasch G, Karim S. The tick endosymbiont Candidatus Midichloria mitochondrii and selenoproteins are essential for the growth of Rickettsia parkeri in the Gulf Coast tick vector. Microbiome. 2018;6:141.

29. Lejal E, Moutailler S, Šimo L, Vayssier-Taussat M, Pollet T. Tick-borne pathogen detection in midgut and salivary glands of adult Ixodes ricinus. Parasites & Vectors. 2019;12:152.

30. Galan M, Razzauti M, Bard E, Bernard M, Brouat C, Charbonnel N, et al. 16S rRNA Amplicon Sequencing for Epidemiological Surveys of Bacteria in Wildlife. mSystems. 2016;1:e00032–16.

31. Aitchison J, Ho CH. The multivariate Poisson-log normal distribution. Biometrika. 1989;76:643–53.

32. Chiquet J, Mariadassou M, Robin S. PLNmodels: Poisson Lognormal Models [Internet]. 2019 [cited 2019 Oct 19]. Available from: https://CRAN.R-project.org/package=PLNmodels

33. Chiquet J, Mariadassou M, Robin S. Variational inference for sparse network reconstruction from count data. arXiv:180603120 [stat] [Internet]. 2018 [cited 2019 Oct 18]; Available from: http://arxiv.org/abs/1806.03120

34. Chiquet J, Mariadassou M, Robin S. Variational inference for probabilistic Poisson PCA. Ann Appl Stat. 2018;12:2674–98.

35. Robinson MD, McCarthy DJ, Smyth GK. edgeR: a Bioconductor package for differential expression analysis of digital gene expression data. Bioinformatics. Oxford Academic; 2010;26:139–40.

36. Robinson MD, Oshlack A. A scaling normalization method for differential expression analysis of RNA-seq data. Genome Biology. 2010;11:R25.

37. McCarthy DJ, Chen Y, Smyth GK. Differential expression analysis of multifactor RNA-Seq experiments with respect to biological variation. Nucleic Acids Res. Oxford Academic; 2012;40:4288–97.

38. R Core Team. R: A Language and Environment for Statistical Computing [Internet]. Vienna, Austria: R Foundation for Statistical Computing; 2020. Available from: https://www.R-project.org/

39. Plantard O, Bouju-Albert A, Malard M-A, Hermouet A, Capron G, Verheyden H. Detection of Wolbachia in the Tick Ixodes ricinus is Due to the Presence of the Hymenoptera Endoparasitoid Ixodiphagus hookeri. PLOS ONE. 2012;7:e30692.

40. Bohacsova M, Mediannikov O, Kazimirova M, Raoult D, Sekeyova Z. Arsenophonus nasoniae and Rickettsiae Infection of Ixodes ricinus Due to Parasitic Wasp Ixodiphagus hookeri. PLOS ONE. 2016;11:e0149950.

41. Heise SR, Elshahed MS, Little SE. Bacterial Diversity in Amblyomma americanum (Acari: Ixodidae) With a Focus on Members of the Genus Rickettsia. J Med Entomol. 2010;47:258–68.

42. Zolnik CP, Prill RJ, Falco RC, Daniels TJ, Kolokotronis S-O. Microbiome changes through ontogeny of a tick pathogen vector. Molecular Ecology. 2016;25:4963–77.

43. Menchaca AC, Visi DK, Strey OF, Teel PD, Kalinowski K, Allen MS, et al. Preliminary Assessment of Microbiome Changes Following Blood-Feeding and Survivorship in the Amblyomma americanum Nymph-to-Adult Transition using Semiconductor Sequencing. PLOS ONE. 2013;8:e67129.

44. Trout Fryxell RT, DeBruyn JM. The Microbiome of Ehrlichia-Infected and Uninfected Lone Star Ticks (Amblyomma americanum). Stevenson B, editor. PLOS ONE. 2016;11:e0146651.

45. Gurfield N, Grewal S, Cua LS, Torres PJ, Kelley ST. Endosymbiont interference and microbial diversity of the Pacific coast tick, Dermacentor occidentalis, in San Diego County, California. PeerJ. 2017;5:e3202.

46. Thapa S, Zhang Y, Allen MS. Effects of temperature on bacterial microbiome composition in Ixodes scapularis ticks. MicrobiologyOpen. 2019;8:e00719.

47. Duron O, Binetruy F, Noël V, Cremaschi J, McCoy KD, Arnathau C, et al. Evolutionary changes in symbiont community structure in ticks. Molecular Ecology. 2017;26:2905–21.

48. Beninati T, Lo N, Sacchi L, Genchi C, Noda H, Bandi C. A Novel Alpha-Proteobacterium Resides in the Mitochondria of Ovarian Cells of the Tick Ixodes ricinus. Appl Environ Microbiol. 2004;70:2596–602.

49. Lo N, Beninati T, Sassera D, Bouman E a. P, Santagati S, Gern L, et al. Widespread distribution and high prevalence of an alpha-proteobacterial symbiont in the tick Ixodes ricinus. Environmental Microbiology. 2006;8:1280–7.

50. Sassera D, Beninati T, Bandi C, Bouman EAP, Sacchi L, Fabbi M, et al. ‘Candidatus Midichloria mitochondrii’, an endosymbiont of the tick Ixodes ricinus with a unique intramitochondrial lifestyle. International Journal of Systematic and Evolutionary Microbiology,. 2006;56:2535–40.

51. Olivieri E, Epis S, Castelli M, Varotto Boccazzi I, Romeo C, Desirò A, et al. Tissue tropism and metabolic pathways of Midichloria mitochondrii suggest tissue-specific functions in the symbiosis with Ixodes ricinus. Ticks and Tick-borne Diseases. 2019;10:1070–7.

52. Schicht S, Schnieder T, Strube C. Rickettsia spp. and Coinfections With Other Pathogenic Microorganisms in Hard Ticks From Northern Germany. J Med Entomol. 2012;49:766–71.

53. Raulf M-K, Jordan D, Fingerle V, Strube C. Association of Borrelia and Rickettsia spp. and bacterial loads in Ixodes ricinus ticks. Ticks and Tick-borne Diseases. 2018;9:18–24.

54. Binetruy F, Bailly X, Chevillon C, Martin OY, Bernasconi MV, Duron O. Phylogenetics of the Spiroplasma ixodetis endosymbiont reveals past transfers between ticks and other arthropods. Ticks and Tick-borne Diseases. 2019;10:575–84.

55. Oliver KM, Russell JA, Moran NA, Hunter MS. Facultative bacterial symbionts in aphids confer resistance to parasitic wasps. PNAS. 2003;100:1803–7.

56. Guay J-F, Boudreault S, Michaud D, Cloutier C. Impact of environmental stress on aphid clonal resistance to parasitoids: Role of Hamiltonella defensa bacterial symbiosis in association with a new facultative symbiont of the pea aphid. Journal of Insect Physiology. 2009;55:919–26.

57. Towner K. The Genus Acinetobacter. In: Dworkin M, Falkow S, Rosenberg E, Schleifer K-H, Stackebrandt E, editors. The Prokaryotes: Volume 6: Proteobacteria: Gamma Subclass [Internet]. New York, NY: Springer New York; 2006 [cited 2019 Oct 11]. p. 746–58. Available from: https://doi.org/10.1007/0-387-30746-X_25

58. Falkinham JO. Environmental Sources of Nontuberculous Mycobacteria. Clinics in Chest Medicine. 2015;36:35–41.

59. Kämpfer P, McInroy JA, Glaeser SP. Chryseobacterium rhizoplanae sp. nov., isolated from the rhizoplane environment. Antonie van Leeuwenhoek. 2015;107:533–8.

60. Horn H, Keller A, Hildebrandt U, Kämpfer P, Riederer M, Hentschel U. Draft genome of the Arabidopsis thaliana phyllosphere bacterium, Williamsia sp. ARP1. Standards in Genomic Sciences. 2016;11:8.

61. An S, Berg G. Stenotrophomonas maltophilia. Trends in Microbiology. 2018;26:637–8.

62. Green PN, Ardley JK. Review of the genus Methylobacterium and closely related organisms: a proposal that some Methylobacterium species be reclassified into a new genus, Methylorubrum gen. nov. International Journal of Systematic and Evolutionary Microbiology,. 2018;68:2727–48.

63. Salter SJ, Cox MJ, Turek EM, Calus ST, Cookson WO, Moffatt MF, et al. Reagent and laboratory contamination can critically impact sequence-based microbiome analyses. BMC Biology. 2014;12:87.

64. Glassing A, Dowd SE, Galandiuk S, Davis B, Chiodini RJ. Inherent bacterial DNA contamination of extraction and sequencing reagents may affect interpretation of microbiota in low bacterial biomass samples. Gut Pathogens. 2016;8:24.

65. Weyrich LS, Farrer AG, Eisenhofer R, Arriola LA, Young J, Selway CA, et al. Laboratory contamination over time during low-biomass sample analysis. Molecular Ecology Resources. 2019;19:982–96.

66. Binetruy F, Dupraz M, Buysse M, Duron O. Surface sterilization methods impact measures of internal microbial diversity in ticks. Parasites Vectors. 2019;12:268.

67. Baldrian P. Microbial activity and the dynamics of ecosystem processes in forest soils. Current Opinion in Microbiology. 2017;37:128–34.

68. Narasimhan S, Fikrig E. Tick microbiome: the force within. Trends in Parasitology. 2015;31:315–23.

69. Narasimhan S, Rajeevan N, Liu L, Zhao YO, Heisig J, Pan J, et al. Gut Microbiota of the Tick Vector Ixodes scapularis Modulate Colonization of the Lyme Disease Spirochete. Cell Host & Microbe. 2014;15:58–71.

70. Landesman WJ, Mulder K, Allan BF, Bashor LA, Keesing F, LoGiudice K, et al. Potential effects of blood meal host on bacterial community composition in Ixodes scapularis nymphs. Ticks and Tick-borne Diseases. 2019;10:523–7.

71. Randolph SE, Green RM, Hoodless AN, Peacey MF. An empirical quantitative framework for the seasonal population dynamics of the tick Ixodesricinus. International Journal for Parasitology. 2002;32:979–89.

72. Abdullah S, Davies S, Wall R. Spectrophotometric analysis of lipid used to examine the phenology of the tick Ixodes ricinus. Parasites & Vectors. 2018;11:523.

73. Eiler A, Heinrich F, Bertilsson S. Coherent dynamics and association networks among lake bacterioplankton taxa. ISME J. 2012;6:330–42.

74. Chow C-ET, Kim DY, Sachdeva R, Caron DA, Fuhrman JA. Top-down controls on bacterial community structure: microbial network analysis of bacteria, T4-like viruses and protists. ISME J. 2014;8:816–29.

75. Fuhrman JA, Steele JA. Community structure of marine bacterioplankton: patterns, networks, and relationships to function. Aquatic Microbial Ecology. 2008;53:69–81.

76. Berdjeb L, Parada A, Needham DM, Fuhrman JA. Short-term dynamics and interactions of marine protist communities during the spring–summer transition. ISME J. 2018;12:1907–17.

77. Abraham NM, Liu L, Jutras BL, Yadav AK, Narasimhan S, Gopalakrishnan V, et al. Pathogen-mediated manipulation of arthropod microbiota to promote infection. PNAS. 2017;114:E781–90.

78. Tully JG, Rose DL, Yunker CE, Carle P, Bové JM, Williamson DL, et al. Spiroplasma ixodetis sp. nov., a New Species from Ixodes pacificus Ticks Collected in Oregon. International Journal of Systematic and Evolutionary Microbiology,. 1995;45:23–8.

79. Henning K, Greiner-Fischer S, Hotzel H, Ebsen M, Theegarten D. Isolation of Spiroplasma sp. from an Ixodes tick. International Journal of Medical Microbiology. 2006;296:157–61.

80. Weinert LA, Tinsley MC, Temperley M, Jiggins FM. Are we underestimating the diversity and incidence of insect bacterial symbionts? A case study in ladybird beetles. Biology Letters. 2007;3:678–81.

81. Duron O, Bouchon D, Boutin S, Bellamy L, Zhou L, Engelstädter J, et al. The diversity of reproductive parasites among arthropods: Wolbachia do not walk alone. BMC Biology. 2008;6:27.

82. Hurst GDD, Schulenburg JHG von der, Majerus TMO, Bertrand D, Zakharov IA, Baungaard J, et al. Invasion of one insect species, Adalia bipunctata, by two different male-killing bacteria. Insect Molecular Biology. 1999;8:133–9.

83. Majerus TMO, Schulenburg JHGVD, Majerus MEN, Hurst GDD. Molecular identification of a male-killing agent in the ladybird Harmonia axyridis (Pallas) (Coleoptera: Coccinellidae). Insect Molecular Biology. 1999;8:551–5.

84. Majerus MEN, Hinrich J, Schulenburg GVD, Zakharov IA. Multiple causes of male-killing in a single sample of the two-spot ladybird, Adalia bipunctata (Coleoptera: Coccinellidae) from Moscow. Heredity. 2000;84:605–9.

85. Jiggins FM, Hurst GDD, Jiggins CD, Schulenburg JHG v d, Majerus MEN. The butterfly Danaus chrysippus is infected by a male-killing Spiroplasma bacterium. Parasitology. 2000;120:439–46.

86. Tinsley MC, Majerus MEN. A new male-killing parasitism: Spiroplasma bacteria infect the ladybird beetle Anisosticta novemdecimpunctata (Coleoptera: Coccinellidae). Parasitology. 2006;132:757–65.

87. Sanada-Morimura S, Matsumura M, Noda H. Male Killing Caused by a Spiroplasma Symbiont in the Small Brown Planthopper, Laodelphax striatellus. J Hered. 2013;104:821–9.

88. Xie J, Butler S, Sanchez G, Mateos M. Male killing Spiroplasma protects Drosophila melanogaster against two parasitoid wasps. Heredity. 2014;112:399–408.

89. Peix A, Ramírez-Bahena M-H, Velázquez E. Historical evolution and current status of the taxonomy of genus Pseudomonas. Infection, Genetics and Evolution. 2009;9:1132–47.

90. Meikle WG, Bon M-C, Cook SC, Gracia C, Jaronski ST. Two strains of Pseudomonas fluorescens bacteria differentially affect survivorship of waxworm (Galleria mellonella) larvae exposed to an arthropod fungal pathogen, Beauveria bassiana. Biocontrol Science and Technology. Taylor & Francis; 2013;23:220–33.

91. Meikle WG, Mercadier G, Guermache F, Bon M-C. Pseudomonas contamination of a fungus-based biopesticide: Implications for honey bee (Hymenoptera: Apidae) health and Varroa mite (Acari: Varroidae) control. Biological Control. 2012;60:312–20.

92. Vodovar N, Vinals M, Liehl P, Basset A, Degrouard J, Spellman P, et al. Drosophila host defense after oral infection by an entomopathogenic Pseudomonas species. PNAS. National Academy of Sciences; 2005;102:11414–9.

93. Vodovar N, Vallenet D, Cruveiller S, Rouy Z, Barbe V, Acosta C, et al. Complete genome sequence of the entomopathogenic and metabolically versatile soil bacterium Pseudomonas entomophila. Nature Biotechnology. Nature Publishing Group; 2006;24:673–9.

94. Evtushenko LI, Takeuchi M. The Family Microbacteriaceae. In: Dworkin M, Falkow S, Rosenberg E, Schleifer K-H, Stackebrandt E, editors. The Prokaryotes: Volume 3: Archaea Bacteria: Firmicutes, Actinomycetes [Internet]. New York, NY: Springer New York; 2006 [cited 2019 Oct 19]. p. 1020–98. Available from: https://doi.org/10.1007/0-387-30743-5_43

95. Jiang Y, Wiese J, Tang SK, Xu LH, Imhoff JF, Jiang CL. Actinomycetospora chiangmaiensis gen. nov., sp. nov., a new member of the family Pseudonocardiaceae. International Journal of Systematic and Evolutionary Microbiology. 2008;58:408–13.

96. Lee SD. Kineococcus rhizosphaerae sp. nov., isolated from rhizosphere soil. International Journal of Systematic and Evolutionary Microbiology,. 2009;59:2204–7.

97. Yamamura H, Ashizawa H, Nakagawa Y, Hamada M, Ishida Y, Otoguro M, et al. Actinomycetospora rishiriensis sp. nov., isolated from a lichen. International Journal of Systematic and Evolutionary Microbiology,. 2011;61:2621–5.

98. Eberl L, Vandamme P. Members of the genus Burkholderia: good and bad guys. F1000Res [Internet]. 2016 [cited 2019 Oct 19];5. Available from: https://www.ncbi.nlm.nih.gov/pmc/articles/PMC4882756/

99. Viana AT, Caetano T, Covas C, Santos T, Mendo S. Environmental superbugs: The case study of Pedobacter spp. Environmental Pollution. 2018;241:1048–55.

100. Burgdorfer W, Hayes SF, Mavros AJ. Nonpathogenic rickettsiae in Dermacentor andersoni: a limiting factor for the distribution of Rickettsia rickettsii. Rickettsiae and Rickettsial Diseases. 1981;585–594.

101. Steiner FE, Pinger RR, Vann CN, Grindle N, Civitello D, Clay K, et al. Infection and Co-infection Rates of Anaplasma phagocytophilum Variants, Babesia spp., Borrelia burgdorferi, and the Rickettsial Endosymbiont in Ixodes scapularis (Acari: Ixodidae) from Sites in Indiana, Maine, Pennsylvania, and Wisconsin. J Med Entomol. 2008;45:289–97.

102. Telford SR. Status of the “East side hypothesis” (transovarial interference) twenty five years later. Annals of the New York Academy of Sciences. 2009;1166:144.

103. Williams-Newkirk AJ, Rowe LA, Mixson-Hayden TR, Dasch GA. Characterization of the Bacterial Communities of Life Stages of Free Living Lone Star Ticks (Amblyomma americanum). Munderloh UG, editor. PLoS ONE. 2014;9:e102130.

104. Tijsse-Klasen E, Sprong H, Pandak N. Co-infection of Borrelia burgdorferi sensu lato and Rickettsia species in ticks and in an erythema migrans patient. Parasites & Vectors. 2013;6:347.

105. Moniuszko A, Dunaj J, Święcicka I, Zambrowski G, Chmielewska-Badora J, Żukiewicz-Sobczak W, et al. Co-infections with Borrelia species, Anaplasma phagocytophilum and Babesia spp. in patients with tick-borne encephalitis. Eur J Clin Microbiol Infect Dis. 2014;33:1835–41.

106. Hoversten K, Bartlett MA. Diagnosis of a tick-borne coinfection in a patient with persistent symptoms following treatment for Lyme disease. BMJ Case Rep [Internet]. 2018 [cited 2018 Nov 19];2018. Available from: http://europepmc.org/abstract/med/30262525

107. Krause PJ, Telford SR, Spielman A, Sikand V, Ryan R, Christianson D, et al. Concurrent Lyme Disease and Babesiosis: Evidence for Increased Severity and Duration of Illness. JAMA. 1996;275:1657–60.

108. Diuk-Wasser MA, Vannier E, Krause PJ. Coinfection by Ixodes Tick-Borne Pathogens: Ecological, Epidemiological, and Clinical Consequences. Trends in Parasitology. 2016;32:30–42.

109. Adegoke A, Kumar D, Bobo C, Rashid MI, Durrani AZ, Sajid MS, et al. Tick-Borne Pathogens Shape the Native Microbiome Within Tick Vectors. Microorganisms. Multidisciplinary Digital Publishing Institute; 2020;8:1299.

